# Generative probabilistic biological sequence models that account for mutational variability

**DOI:** 10.1101/2020.07.31.231381

**Authors:** Eli N. Weinstein, Debora S. Marks

## Abstract

Large-scale sequencing has revealed extraordinary diversity among biological sequences, produced over the course of evolution and within the lifetime of individual organisms. Existing methods for building statistical models of sequences often pre-process the data using multiple sequence alignment, an unreliable approach for many genetic elements (antibodies, disordered proteins, etc.) that is subject to fundamental statistical pathologies. Here we introduce a structured emission distribution (the MuE distribution) that accounts for mutational variability (substitutions and indels) and use it to construct generative and predictive hierarchical Bayesian models (H-MuE models). Our framework enables the application of arbitrary continuous-space vector models (e.g. linear regression, factor models, image neural-networks) to unaligned sequence data. Theoretically, we show that the MuE generalizes classic probabilistic alignment models. Empirically, we show that H-MuE models can infer latent representations and features for immune repertoires, predict functional unobserved members of disordered protein families, and forecast the future evolution of pathogens.

## 1 Introduction

High-throughput sequencing has become pervasive across modern biology and biomedicine, and has revealed extraordinary sequence diversity among proteins, RNA, and other genetic elements. Interpreting that diversity, and making predictions about unobserved or future sequences, is an open challenge, with relevance to epidemiology (predicting pathogen evolution), immunology (characterizing antibody repertoires), molecular evolution (mapping substructure within protein families), protein design, and far more. Accomplishing these goals requires tools for working with high-dimensional complex sequence distributions. In principal, generative probabilistic models of biological sequences could enable discovery of rare subpopulations, key sequence features, trends across time, the impact of experimental interventions, etc., and then convert this understanding into predictions of new sequences that could be synthesized and tested in the laboratory.

There are a variety of methods that have seen widespread success on similar challenges in other fields of science, but adapting them to structured data such as biological sequences is non-trivial. In particular, there is an enormous wealth of generative models of continuous-space vectors, such as linear regression, probabilistic PCA [1], non-negative matrix factorization [2], and image neural networks [3]. In order to apply continuous-space vector models to types of data besides continuous-space vectors, statisticians typically rely on an *emission* distribution, e.g. a Poisson distribution for rare count data or a zero-inflated negative binomial distribution for single cell RNA expression data [4, 5, 6]. For instance, if *p*(*v*|*θ*) is a generative model of continuous-space vectors, with parameter *θ*, a generative model of count vectors *y* can be built with a Poisson emission distribution as:

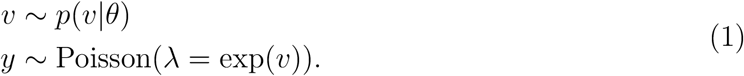

While there is often a wide variety of possible choices for an emission distribution, a good emission distribution should not only generate the right type of data, but also capture the variability commonly seen in the data.

To enable the principled application of continuous-space vector models to biological sequences, we propose the “mutational emission”, or “MuE” distribution. This new distribution generates unaligned sequence data while explicitly accounting for the kinds of variability commonly seen in biological sequence data, namely substitution, insertion and deletion mutations. Using the MuE as an emission distribution enables the direct application of continuous-space vector models to biological sequences; we term these combined models “hierarchical MuE” or H-MuE models. H-MuE models do not require a multiple sequence alignment of the data, which is often unreliable in practice, especially for disordered proteins, antibodies, promoters, and other genetic elements, and is pathological in theory when the ultimate aim is sequence prediction (Section S2). Instead, H-MuE models represent the multiple sequence alignment implicitly as a latent variable, making it possible to account rigorously for alignment uncertainty. Classical probabilistic alignment models can be re-derived as special cases of the MuE distribution. Moreover, in contrast to alternative models of biological sequence mutation, the likelihood function of the MuE distribution is analytically tractable and differentiable [7]. This helps enable inference of the parameters of H-MuE models from data using Bayes’ rule, allowing in particular the use of scalable approximate inference algorithms that rely on automatic differentiation (also known as backpropagation) [8, 9].

We demonstrate empirically how H-MuE models can be applied to large sequence datasets to map the biological diversity found among disordered proteins, viruses, immune receptors, and more. We show that H-MuE models can be used to predict unobserved and future protein sequences, and enable unsupervised learning of sequence subpopulations and sequence features.

## 2 Results

### 2.1 The MuE distribution

We consider datasets of unaligned biological sequences *y*_*i*_ for *i* ∈ {1, …, *N*}, which may be recorded from different species, organisms, cells, etc. In order to model the distribution of sequences, and how this distribution may depend on any covariates, we designed two-layer generative models. The first layer generates an “ancestral” sequence logo *x*_*i*_ with fixed length; the second step adds mutations, including substitutions, insertions and deletions to generate *y*_*i*_. The first step relies on a continuous-space matrix model *p*(*v*|*θ*), where *v* ∈ ℝ^*M*×*D*^, which may be any model of interest and may depend on covariates such as sequence collection time. The second step employs the proposed mutational emission distribution MuE(*x, c, a, ℓ*) (details in Box 1), which describes a distribution of mutants of *x* with indel probabilities controlled by the parameter *a*, substitution probabilities controlled by the parameter *ℓ*, and insertion sequences controlled by the parameter *c* (Figure 1A). The complete H-MuE model is:

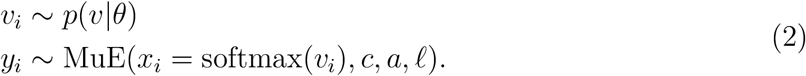

where the softmax linker function sets *x*_*i,m,d*_ = exp(*v*_*i,m,d*_)/ ∑_*d*′_ exp(*v*_*i,m,d*′_) for *m* ∈ {1, …, *M*} and *d, d′* ∈ {1, …, *D*}. We will refer to models that fit the form of Equation 2 as “hierarchical MuE” (H-MuE) models, since they generate sequences according to a hierarchical Bayesian model, first employing *p*(*v*|*θ*) to generate ancestral sequence logos *x*_1_, …, *x*_*N*_, then employing the MuE distribution to add additional mutations (Figure 1B). More generally, *c*, *a* and *ℓ* may also be made to depend on *v*_*i*_ (Section S5).

**Figure 1:**
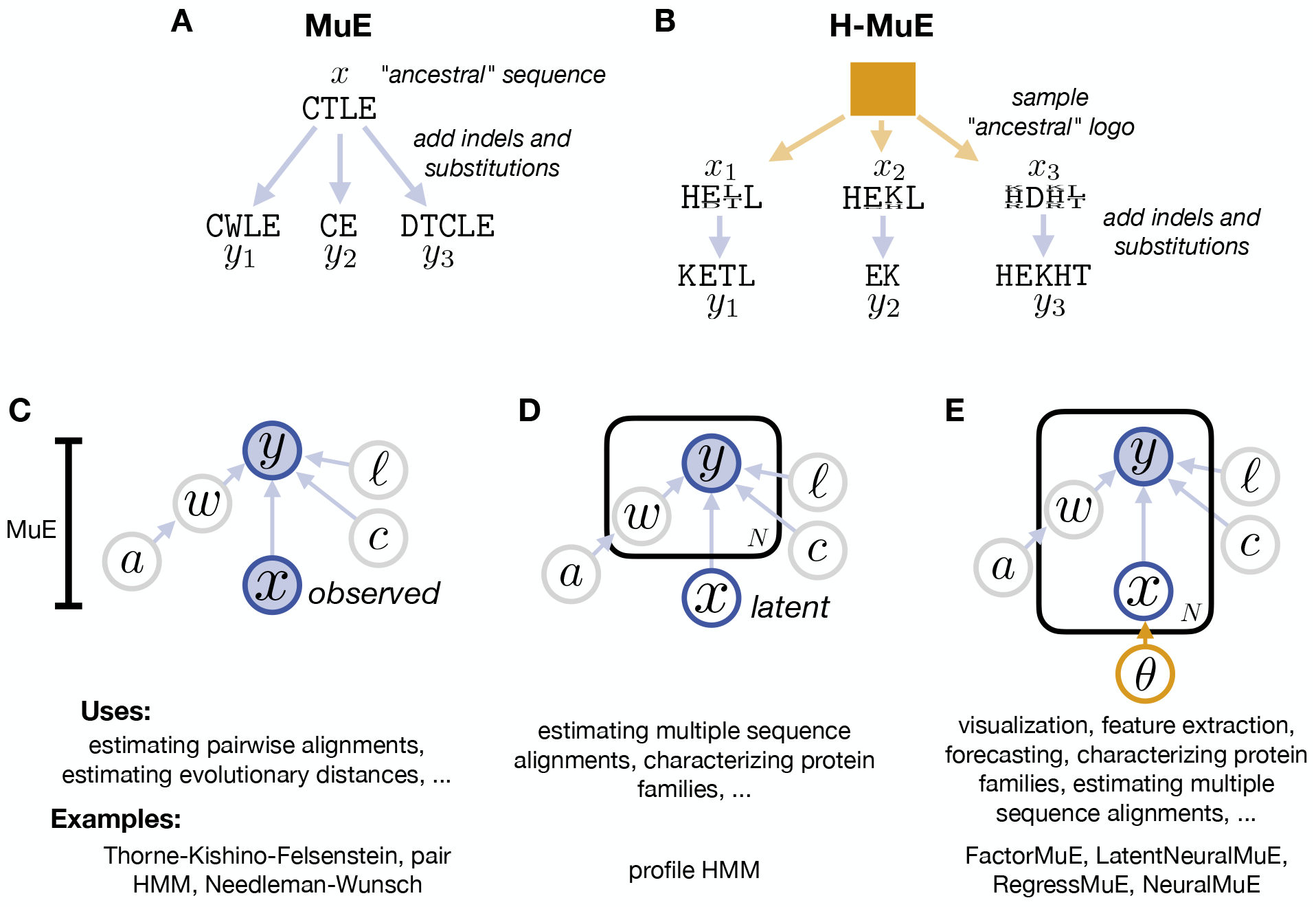
Hierarchical mutational emission (H-MuE) models generalize and extend previous probabilistic mutation models and alignment methods. A. The MuE model generates samples *y*_*i*_ that are mutants of an initial sequence *x* (see Figure S1 for a detailed illustration). B. In an H-MuE model, *v*_*i*_ is sampled from an initial continuous vector model *p*(*v*|*θ*) and determines the sequence logo *x*_*i*_ = softmax(*v*_*i*_); then *y*_*i*_ is sampled from *x*_*i*_ according to the MuE distribution (see Figure S2 for a detailed illustration). C,D,E. Graphical models of alternative use cases for the MuE distribution, along with examples of models for each use case (based on our theoretical results, Section S4). C. In some situations, sequence *x* is observed as well as *y*; estimating the hidden state variable of the MuE distribution can provide an alignment between the two sequences, and estimating the parameters can give insight into their evolutionary relatedness. D. A collection of sequences *y*_1_, …, *y*_*N*_ can be modeled as mutants of an unobserved “ancestral” sequence or sequence logo. The pHMM fits this graphical model. E. In H-MuE models, each sequence *y*_*i*_ is associated with its own individual “ancestor” *x*_*i*_, drawn from a population determined by *θ*.

Given a dataset of sequences *y*_1:*N*_, we would like to infer the parameters of an H-MuE model (in particular, *θ*, *c*, *a* and *ℓ*) using Bayes’ theorem. While exact inference is computationally intractable, we can approximate the posterior distribution using variational inference [10]. We use stochastic black box variational inference with the reparameterization trick, relying on the crucial property of the MuE distribution that its likelihood *p*_MuE_(*y_i_|x_i_, c, a*, *ℓ*) is a differentiable function of the parameters *x*_*i*_, *c*, *a* and *ℓ* [11]. For models with local latent variables, we also use a recognition network to amortize computation across datapoints [12, 13]. When implemented for modern computing hardware (graphics processing units), these techniques together enable scalable approximate inference in H-MuE models for a wide variety of continuous-space models *p*(*v*|*θ*) (Section S6) [8].

##### Box 1: MuE Mathematical Details

The MuE is a structured hidden Markov model (HMM), with initial transition vector *a*^(*i*)^, transition matrix *a*^(*t*)^ and discrete emission matrix *e* = (ξ · *x* + ζ · *c*) · *ℓ*, where ξ and ζ are fixed constants. The matrix ξ has shape *K × M*, *x* has shape *M × D*, ζ has shape *K* × (*M* +1), *c* has shape (*M* +1) × *D* and *ℓ* has shape *D × B*, where *M* is the length of the ancestral sequence, *K* = 2(*M* + 1) is the size of the Markov chain state space, *D* is the alphabet size of the ancestral sequence and *B* is the alphabet size of the observed sequence (typically *D* = *B*). Each row of the matrices *x*, *c* and *ℓ* is a vector of discrete probabilities that sum to one, ie. 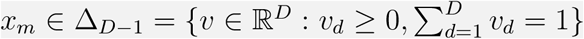. The constants ξ and ζ are defined by ξ_*k,m*_ = δ_*k*,2*m*_ and ζ_*k,m*_ = δ_*k*,2*m*−1_, where δ_*k,k′*_ is the Kronecker delta, ie. δ_*k,k′*_ = 1 if *k* = *k′* and δ_*k,k′*_ = 0 if *k* ∕= *k′*. Crucially, the transition matrix *a*^(*t*)^ must satisfy the restriction that 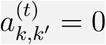 for all *k, k′* such that (1) state *k* is accessible from the initial state and (2) *k′* + *k′*%2 − *k* + *k*%2 ≤ 0, where % is the modulo (remainder) operation.

To see how the MuE represents alignments, we rewrite MuE(*x, c, a*, *ℓ*) as

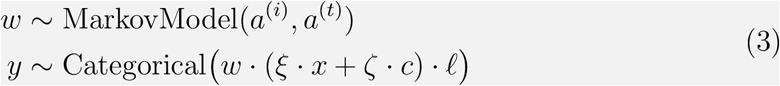

where *w* is a one-hot encoding of the states of a trajectory sampled from the Markov model. *w* has size *L × K*, where *L*, the length of the trajectory, is itself a random variable and determines the length of the sampled sequence *y* (Section S3.1). We see from Equation 3 that *w* acts like a matrix of regression coefficients, determining which position in the regressor ancestral sequence *x* influences each position in the regressand mutated sequence *y*. In Section S4.1 we formalize a mapping between the variable *w* and a pairwise alignment between *x* and *y*, and prove that the restrictions on the transition matrix *a*^(*t*)^ are necessary and sufficient to guarantee that this mapping exists.

### 2.2 The MuE distribution generalizes probabilistic sequence alignment models and methods

Before employing H-MuE models in practice, we analyzed the MuE distribution theoretically. First we showed that if we choose the parameters *a* and *ℓ* such that there is zero probability of insertions, deletions or substitutions, the MuE distribution reduces to a multivariate categorical distribution, a standard emission distribution for pre-aligned sequences (Section S3.2). We used this fact to show that H-MuE models generalize state-of-the-art aligned sequence models [14].

Next, we showed that standard probabilistic mutation and alignment models are also special cases of the MuE distribution. First, we proved that for a particular setting of *a* and *ℓ*, the MuE distribution reduces to the transition distribution of the Thorne-Kishino-Felsenstein model, a continuous-time stochastic process model of sequence evolution that includes indels and satisfies detailed balance (Section S4.2) [15]. Second, we showed that for another setting of *a* and *ℓ*, the MuE distribution matches the conditional distribution of *y* given *x* under the pair hidden Markov model, a common probabilistic pairwise sequence alignment model (Section S4.3) [16]. These two models have usually been employed in contexts where *x* is observed (that is, *x* is a one-hot encoding of a particular sequence) (Figure 1C). Next, we showed that for another setting of *a*, and with *ℓ* the identity matrix, the MuE distribution reduces to the profile hidden Markov model (pHMM) (Figure S4.4) [16]. The pHMM has usually been employed in contexts where *x* is latent and we observe many sequences *y*_1:*N*_ (Figure 1D). While none of these previous models have been explicitly used as emission distributions (Figure 1E), they have been highly successful across a range of other biological sequence analysis problems, providing evidence that the MuE model can effectively capture common forms of variability among biological sequences.

Finally, we specifically investigated the connection between the MuE distribution and biological sequence alignments. The MuE distribution is a type of structured hidden Markov model, and, in the context of the previous probabilistic alignment models discussed above, the state variable of the hidden Markov model has been interpreted as an alignment between sequences *x* and *y*. Indeed, in addition to probabilistic alignment methods, the MuE is also connected to non-probabilistic alignment methods: we proved that for yet another setting of *a* and *ℓ* the maximum *a posteriori* estimator of the state variable corresponds to the Needleman-Wunsch alignment between *x* and *y* (Section S4.5). In general, we established necessary and sufficient conditions on the transition matrix of the MuE distribution guaranteeing that the state variable can always be interpreted as an alignment (Section S4.1). As a consequence, any MuE or H-MuE model can be used not only as a generative sequence model but also as a probabilistic sequence alignment method, and Bayesian inference of the H-MuE parameters yields a posterior distribution over multiple sequence alignments of the dataset. Our variational inference procedure for H-MuE models marginalizes out the MuE state variable during training, in effect considering all possible alignments of the dataset to address the problem of alignment uncertainty.

### 2.3 H-MuE models describe complex sequence diversity

#### Prediction

We first sought to evaluate the ability of H-MuE models to accurately model the distribution of sequences within protein families and to generate new sequences that satisfy the implicit functional constraints of those families. As a baseline we compared our models to a profile HMM, the most widely-used generative model of unaligned protein sequence families (Section S5.1). We then used the MuE to extend two of the most successful continuous-space vector models, probabilistic PCA and a neural network latent variable model, to unaligned sequences, creating the “FactorMuE” and the “LatentNeuralMuE” respectively (Sections S5.3 and S5.5). These three models – the pHMM, FactorMuE and LatentNeuralMuE – have sequentially increasing model complexity; the FactorMuE and LatentNeuralMuE are capable of representing long-distance epistasis. We measured the capacity of each model to predict unobserved, presumably functional, members of the protein family (Figure 2A). We quantified performance in terms of the heldout per residue perplexity, a metric that is monotonically related to the heldout log likelihood, and is interpretable as the effective number of amino acid choices per position. Per residue perplexity ranges from 1 (perfect prediction) to 20 (naive prediction); a BLOSUM62 substitution matrix model, if it exactly describes the data, will have per residue perplexity of 11.0 (Section S7). We applied these models to five datasets of protein families, ranging in size from 1,000 to 10,000 sequences (Section S8). Four were taken from non-redundant sequence databases: sequences similar to dihydrofolate reductase (DHFR; a widely conserved enzyme, often studied with generative models of aligned sequences), serine recombinase (PINE; a tool for genomic engineering), cyclin dependent kinase inhibitor 1B (CDKN1B/p27; a cell cycle inhibitor with a disordered region) and the human papillomavirus E6 protein (VE6; an oncogenic protein with a disordered region) [17, 18, 19]. The final dataset consisted of human T-cell receptor (TCR) sequences from a healthy donor obtained using single cell sequencing (Section S9). We evaluated model performance on a randomly held out 10% of sequences. The results, summarized in Figure 2B, show that FactorMuE models offer a consistent and large improvement in predictive power over the standard pHMM model in every dataset, with an average change in perplexity of 1.50 (log Bayes factor > 10^3^ across all datasets). Particularly dramatic improvements are seen in the TCR dataset, where perplexity falls by more than 3 (log Bayes factor > 10^4^). Meanwhile, the more complex LatentNeuralMuE model also improves over the pHMM in each dataset and overall (average perplexity change −0.42), but underperforms relative to the simpler FactorMuE model, illustrating the advantages of using the MuE distribution to explore models of different complexity.

**Figure 2:**
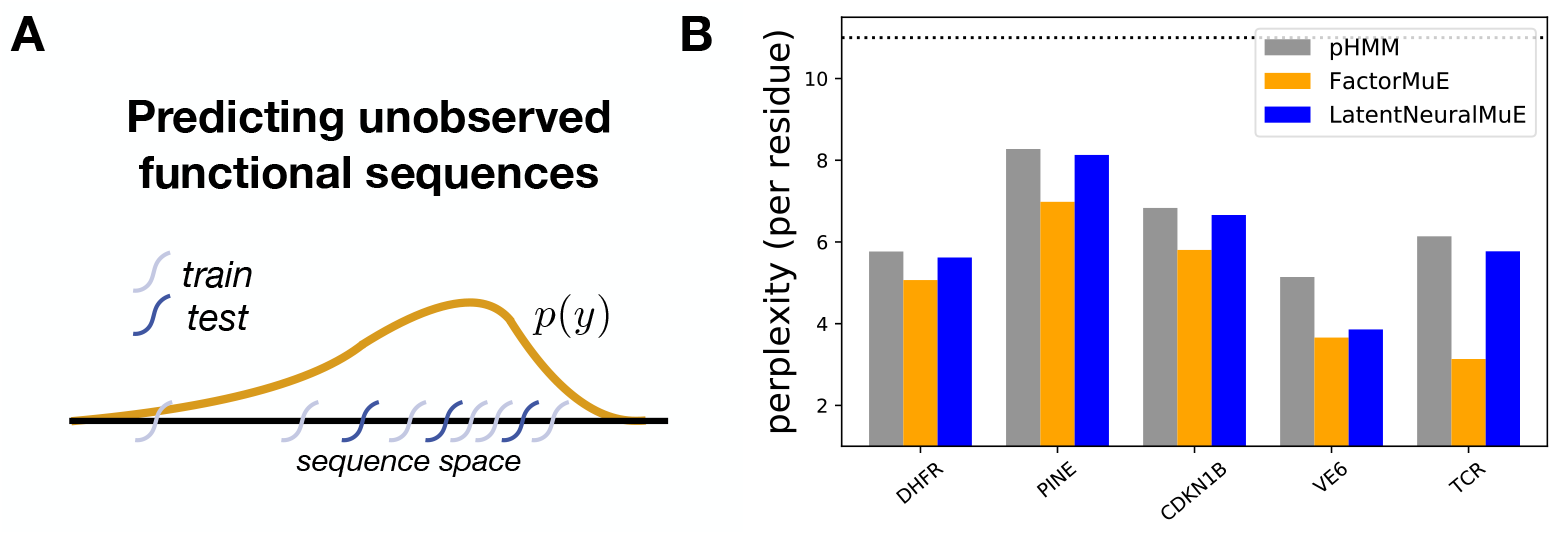
Predicting unobserved functional sequences with H-MuE models. A. Illustration of random train-test data split, used to validate predictions of unobserved members of a sequence family (density estimation). B. Per residue perplexity on the heldout dataset, for the baseline pHMM and two H-MuE models, across five sequence datasets; lower numbers indicate better performance. A BLOSUM62 substitution model that exactly describes the data distribution will have perplexity of 11.0, indicated by a dotted line (Section S7).

#### Latent space

Continuous-space vector models like probabilistic PCA are widely used in other fields to produce visualizations of complex datasets. We examined the two-dimensional embeddings produced by the FactorMuE model, which combines probabilistic PCA with the MuE emission distribution (Figure 3A). We focused on the TCR dataset, and evaluated the model’s capacity to learn richly structured representations in an unsupervised way by cross referencing the latent representation with supervised annotations of the V, D and J segment types in each sequence. We found that the latent space is divided evenly in two, with one side containing TCR*α* sequences and one side TCR*β* sequences (Figure 3B). Each side contains clusters, which correspond to each type of V segment (Figures 3C and S4B). Meanwhile, J segments are distributed uniformly across their corresponding *α* or *β* half, reflecting their ability to recombine with different V segments (Figures 3D and S4C).

**Figure 3:**
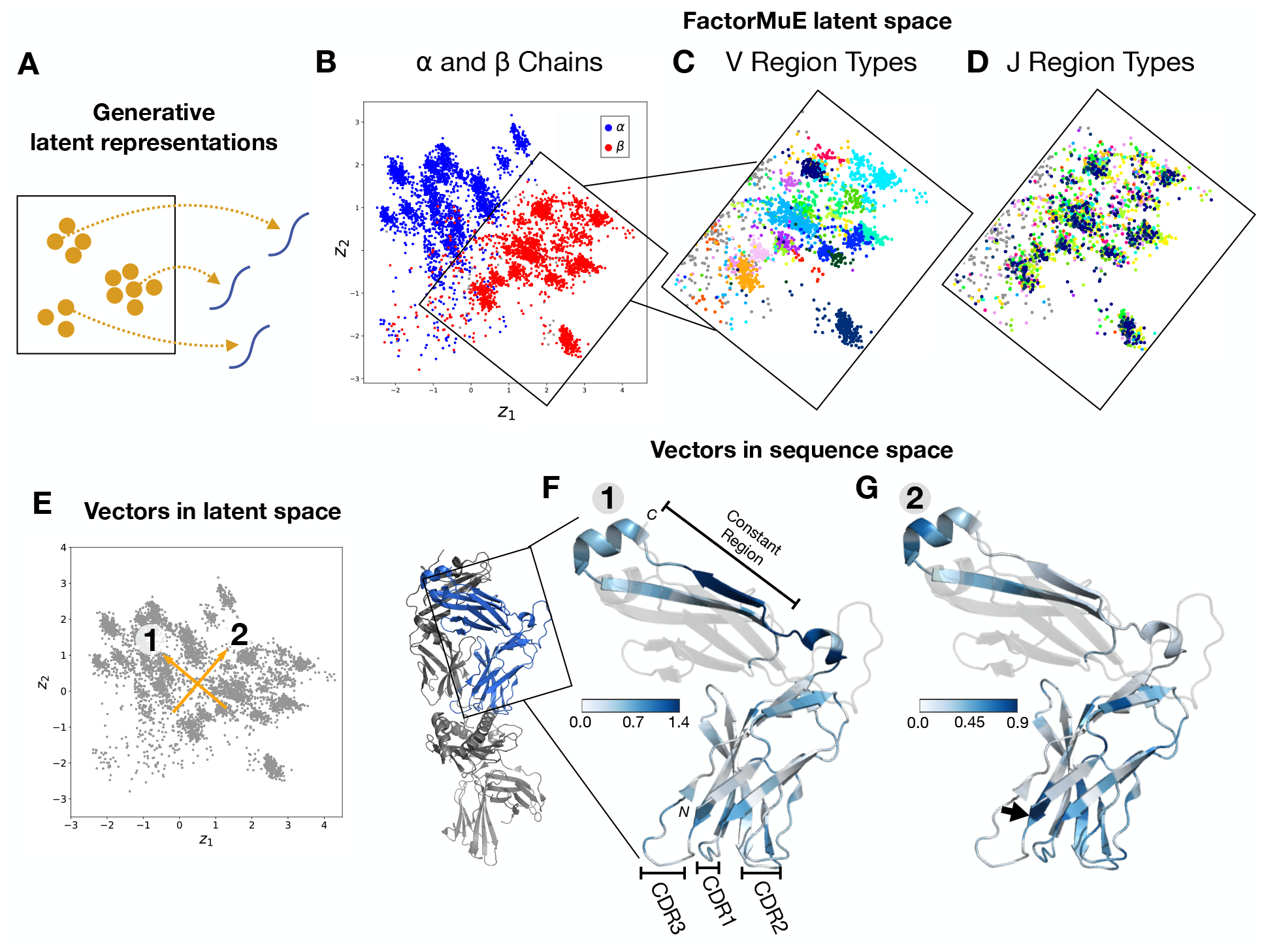
H-MuE model of a single-cell T-cell receptor repertoire. A. Cartoon of a generative model with a latent space sequence representation. B,C,D. The two-dimensional latent space of a FactorMuE model, learned from a TCR dataset taken from a single-cell sequencing experiment. B. Each sequence is colored according to its annotated chain type (grey corresponds to unannotated sequences). C. Each sequence is colored according to its V region type (*α* chains are excluded from the plot for visual clarity, see Figure S4). D. Each sequence is colored according to its J region type. E. A latent space vector normal to the hyperplane separating *α* from *β* chains (vector 1) and an orthogonal vector of the same length (vector 2). F,G. The projection of vectors 1 and 2 back into sequence space, using a reference TCR structure (PDB:2BNR). Residues are colored according to the norm of the shift in amino acid preference from the tail to the head of the latent vector (Equation S78). Transparent residues in the constant region correspond to the portion of the protein that was not sequenced in the experiment. The black arrow in G indicates the start of the CDR3 region, corresponding to the position with the largest shift in preference (Figure S6).

#### Features

By projecting latent space vectors back into sequence space, with the latent MuE alignment variable fixed, we can visualize the features learned by the FactorMuE model and obtain an overview of the major axes of variation in the human TCR repertoire (Section S9). Note that this approach differs fundamentally from standard analysis techniques which focus on cataloguing the usage frequency of different segments or CDR3s in that it describes what the fine-grained variation adds up to at the population level. Consistent with the annotation of the latent representation, the vector normal to the hyperplane separating TCR*α* from TCR*β* chains in the latent space (vector 1) primarily determines the sequence of the constant region of the TCR, while the orthogonal vector (vector 2) primarily determines the sequence of the V chain (Figure 3FG). Along vector 2 we found weak positive correlation between the magnitude of variation and the relative surface accessibility of each site (Spearman correlation *ρ* = 0.20, *p* < 0.02) (Figure S5). The region of largest variation, however, was the buried C-terminal end of the V segment, corresponding to the start of the CDR3 region, the key specificity-determining region (Figure S6). Interestingly, even along vector 1 we observe variation within the variable region, suggesting that there are systematic and heterogeneous differences between the V segment sequence distribution used in TCR*α* chains and that used in TCR*β* chains. To confirm this observation, we developed another H-MuE model: we extended a linear regression model with a MuE distribution (RegressMuE) (Section S5.2). We then used the RegressMuE model to predict the entire TCR sequence based just on its annotation as a TCR*α* or TCR*β*. Figure S7 plots the shift in amino acid preference between the two chains, showing that at a population level there are key positions within the variable region with substantial differences in preference.

### 2.4 H-MuE models forecast sequence evolution

#### Prediction

We evaluated the capacity of H-MuE models to forecast future sequence evolution (Figure 4A). Influenza A is responsible for an estimated 500,000 deaths a year and is an ongoing pandemic threat [20]. It is also a model organism for understanding the dynamics of rapidly evolving pathogens, and forecasting its evolution is essential in preparing vaccines and designing therapeutics [21, 22]. Previous forecasting methods have focused on predicting the relative fitness of existing strains in future years, e.g. [21, 23], or the antigenic properties of newly emerged strains, e.g. [24]. We instead predict the full amino acid sequence of the HA1 protein, the primary site of interaction with the immune system [21, 25]. From the GISAID database we constructed a training set of influenza A(H3N2) HA1 sequences collected from patient samples from 1968 through 2013, and evaluated our predictions on sequences collected from 2014 through October 2019 (420 out of 2,042 sequences held out, 21% of the dataset) (Section S10) [26]. In contrast to the datasets considered in Section 2.3, indels are considered rare, though not absent, in patient samples, and so this dataset also offers an opportunity to evaluate H-MuE models in a distinct regime from that considered previously.

**Figure 4:**
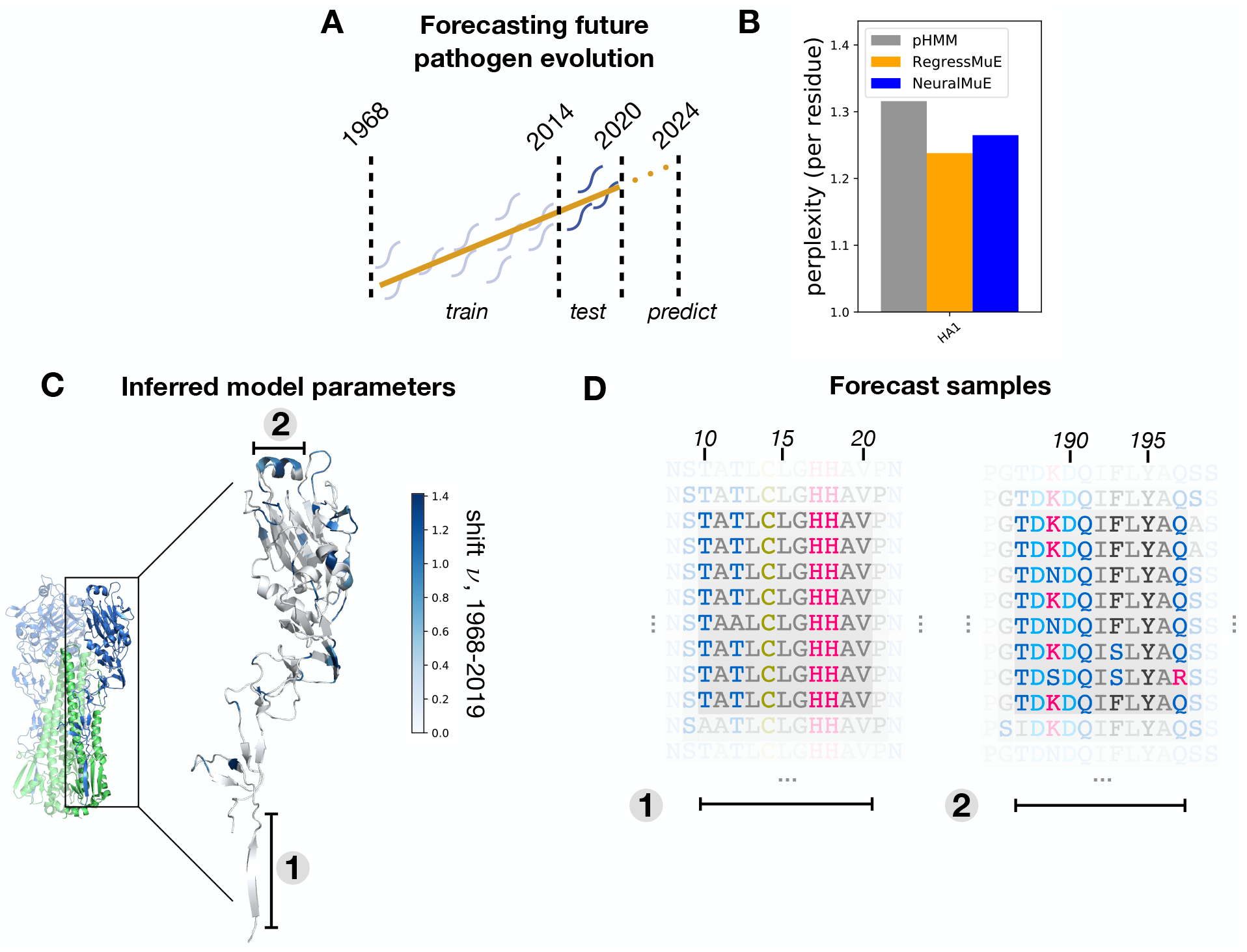
H-MuE model of influenza A(H3N2) evolution. A. Diagram of train-test split for influenza A(H3N2) forecasting. B. Per residue perplexity on the heldout test dataset, for the baseline pHMM and two H-MuE models; lower numbers indicate better performance. C. Magnitude of the shift in amino acid preferences over time inferred by the RegressMuE, *ν*_*l*_ (Equation S80), projected onto an HA1 structure (PDB:4O5N). The full hemagglutinin protein is shown in a smaller size on the left. Region 1 is a portion of the stalk with a small shift over time, while region 2 is the 190 helix, a crucial antigenic region with a particularly large shift over time (Figures S8 and Figure S9). Comparisons with relative solvent accessibility and reproductive fitness measurements are given in Figures S10 and S11. D. Segments of sequences sampled from the posterior predictive distribution for the year 2024. The alignment variable is fixed based on the reference structure (PDB:4O5N), such that segments 1 and 2 correspond to the annotated structural features in C, and the column numbering is standard for influenza A(H3N2) (Section S10).

We considered three different models with increasing complexity. First, the pHMM describes the observed sequences as samples from a population with fixed amino acid frequencies at each site (Section S5.1). The pHMM can capture the observation that there exist key highly variable sites in the HA1 protein, the underlying motivation behind previous prediction methods such as [23]. Next we incorporated sequence collection time as a covariate, using the RegressMuE (Section S5.2). This model, unlike the pHMM, takes into account the possibility that amino acid frequencies may shift over time. Finally, to capture more complex nonlinear dependencies between sequences and time, we extended a neural network regression model with a MuE distribution (NeuralMuE) (Section S5.4). The pHMM achieves a low per residue perplexity of 1.32 but the RegressMuE improves this to 1.24 (log Bayes factor > 10^3^) (Figure 4B). This per residue perplexity difference corresponds to a factor of 10^10^ improvement in per sequence perplexity, a substantial reduction in the space of future viral sequences that must be considered. The NeuralMuE has similar predictive performance to the RegressMuE, with a per residue perplexity of 1.26.

#### Features

Given the success of the RegressMuE in predicting sequences, we investigated in detail what the model can tell us about how HA1 proteins have changed over time. We plotted the magnitude of the shift in amino acid preference from 1968 to 2019 inferred by the model, *ν*_*l*_, for each residue position *l*, with the latent MuE alignment variable kept fixed (Equation S80). We found that sites with large shift are often associated with antigenicity, consistent with the hypothesis that immune evasion is a key driver of influenza evolution. Residues that make up the classical epitope regions A-E of influenza show significantly larger shifts as compared to residues outside these regions (mean *ν*_*l*_ of 0.54 in epitopes A-E versus 0.09 in non-epitope sites, Mann-Whitney U test *p* < 1*e* − 18) (Figure S9) [25, 27]. The same observation holds for residues identified as key determinants of immune escape in recent high-throughput mutational antigenic profiling experiments (mean *ν*_*l*_ of 0.80 in sites with antigenic selection versus 0.24 elsewhere, Mann-Whitney U test *p* < 1*e*−4) (Section S10) [28].

#### Generation

H-MuE models can be used to generate samples of future sequences, enabling experimental tests of immune response and antibody titer on sequences that are likely to emerge in the future. We generated samples for the year 2024 from the RegressMuE and confirmed that they are consistent with previously observed sequences (Figure 4D) (Section S10).

#### Latent space

In stark contrast to the TCR dataset, the latent space representation of the influenza HA1 dataset learned by the FactorMuE model shows the data falling roughly along a line (Figure 5A) (Section S10). The position of a sequence along this line is linearly proportional to the time at which the sequence was collected, though this information was not included in the model (correlation coefficient *ρ* = 0.94) [29]. This observation is consistent with the empirical success of the RegressMuE in sequence prediction, since the RegressMuE model is equivalent to a FactorMuE model with an observed latent representation (Section S5.3). Two clusters of outliers violate the proportionality rule (Figure 5B). The first cluster (marked by ‡) appears around 2008 but the latent representation of these sequences is close to that of sequences from the late 1960s or 1970s; this cluster comes from an experiment performed in 2008 on 1968 sequences, rather than contemporary patient samples as in the rest of the dataset. This observation illustrates how latent representations can be used to clean misannotated sequence data. The second cluster (marked by †) appears in the early 2010s, but the latent representation of these sequences is close to that of sequences from the mid-1990s to early 2000s. Among this cluster of sequences, the ones that have been fully annotated were all collected from an outbreak in the United States of A(H3N2)v triple-reassortant viruses containing matrix protein genes from pandemic A(H1N1)pdm09. In 1998, A(H3N2)- derived viruses jumped from humans to swine, causing a large outbreak among swine, before recombining with other strains to produce this A(H3N2)v outbreak among humans in the 2010s [30, 31]. The epidemiological history is consistent with our unsupervised latent representation, which shows that the cluster of outliers appearing in 2010-2013 most closely matches human samples last seen around 2000.

**Figure 5:**
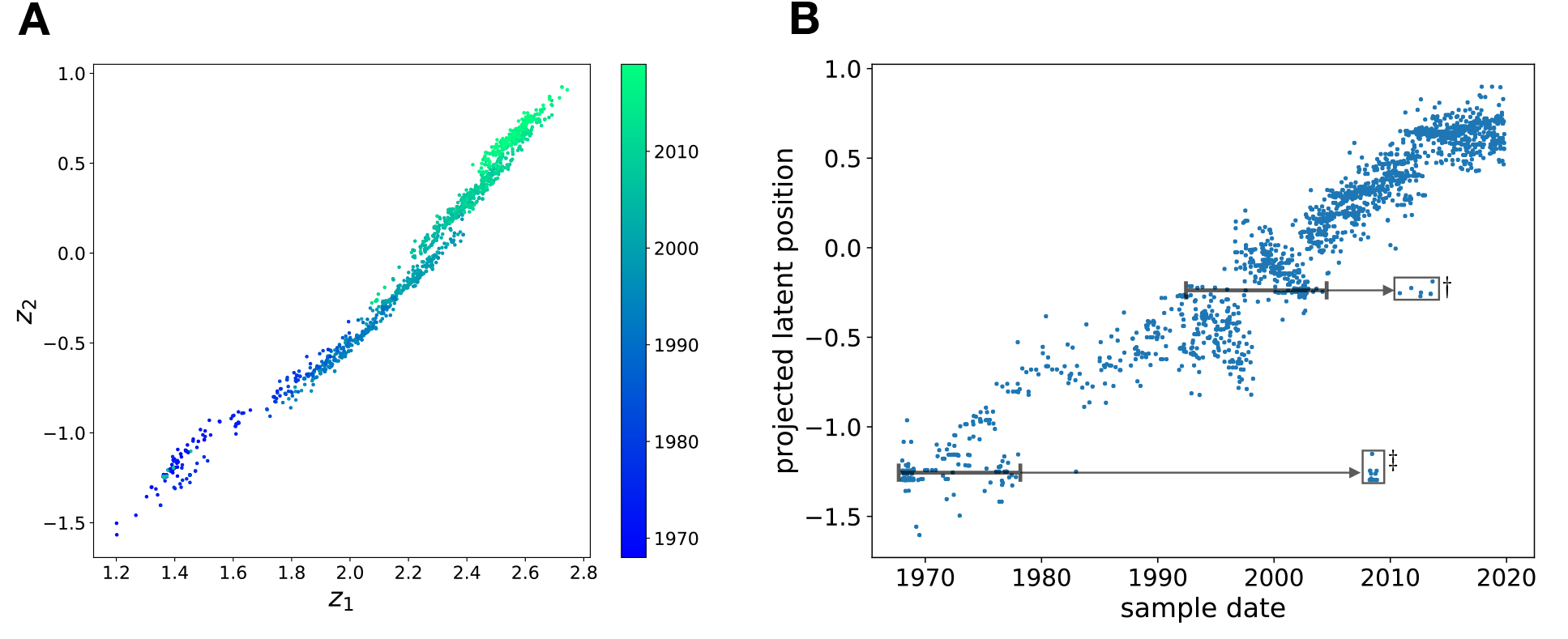
H-MuE model’s latent space representation of influenza A(H3N2) evolution. A. The two-dimensional latent space of the FactorMuE model applied to the A(H3N2) influenza dataset (Section S10). Sequences are colored by the time at which the samples were collected. B. Y-axis: orthogonal projection of the latent representation of each sequence onto the least squares fit line relating *z*_1_ and *z*_2_. X-axis: time at which each sample was collected. Two clusters of outliers are marked († and ‡), along with the time period at which sequences with similar latent representations were last seen (brackets).

## 3 Discussion

H-MuE models are related to previous and contemporaneous work on hierarchical HMM models, such as methods that combine neural networks with HMMs [32] and Potts models with pHMMs [33]. Our work goes further by (1) offering a unified and comprehensive framework for HMM-based probabilistic alignment methods, (2) using probabilistic alignment models as general-purpose emission distributions (and showing that the conventional categorical emission distribution represents a special case), and (3) providing general and scalable approximate Bayesian inference algorithms. To assist in the creation of new models and methods, we have made available an implementation^1^ of the MuE distribution within the probabilistic programming language Edward2, enabling rapid development and testing of new H-MuE models [34].

The MuE distribution is a widely applicable tool for building generative and predictive probabilistic models of biological sequences. Using the MuE distribution, we can extend arbitrary continuous-space vector models to become H-MuE models of unaligned biological sequences, while accounting for mutational variability and statistical uncertainty. Since H-MuE models avoid the pathologies of MSA-based methods, we can apply H-MuE models in a wide variety of statistical contexts, including causal inference, semi-supervised learning, decision-making and design problems. In this article we have explored common vector models (linear regression, probabilistic PCA, a regression neural network and a neural network latent variable model) and focused on prediction and forecasting problems. We anticipate that a wide variety of other probabilistic models and methods – such as nonlinear time series models, sparse regression models, and independent component analysis models – could have as great an impact on biological sequence statistics as they have had on other fields.

## Acknowledgments

We wish to thank Chris Sander, John Ingraham, Elizabeth Wood, and members of the Marks lab for discussion and suggestions. E.N.W. is supported by the Fannie and John Hertz Foundation. D.S.M is supported by the Chan Zuckerberg Initiative.

## Supplementary material

### S1 Supplementary Figures

**Figure S1:**
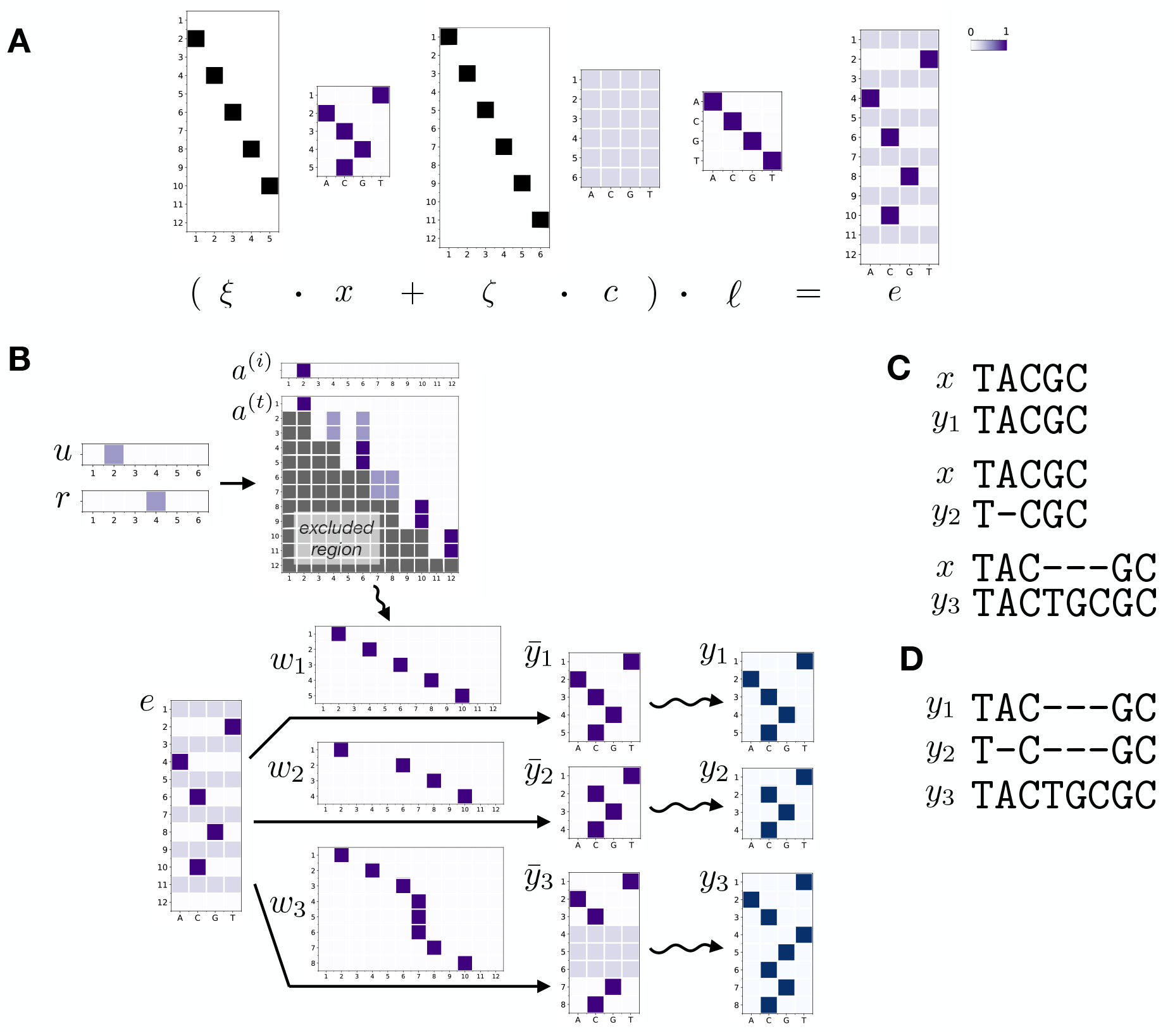
Detailed example of the MuE distribution. A. Constructing the emission matrix *e* = (ξ · *x* + ζ · *c*) · *ℓ* for example *x*, *c* and *ℓ* matrices. The “ancestral” DNA sequence is set to `TACGC`; the matrix *x* is a one-hot encoding of this sequence. B. Illustration of three independent samples *y*_1_, *y*_2_ and *y*_3_ from MuE(*x, c, a*, *ℓ*). Straight lines show deterministic dependencies between variables; squiggly lines show stochastic dependencies. In this example, the transition matrix *a*^(*t*)^ and initial transition vector *a*^(*i*)^ depend on the parameters *u* and *r* as given in Section S5. In grey is the excluded region of the transition matrix which must be zero, ie. where *k′* + *k′*%2 − *k* + *k*%2 ≤ 0. Each state trajectory *w*_*i*_ is sampled independently from a Markov model with the given transition matrix. We also show 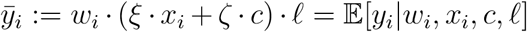. In this example, the generated mutant sequences *y*_1_, *y*_2_ and *y*_3_ are TACGC, TCGC and TACTGCGC. C. Pairwise alignments between *x* and each *y*_*i*_, as determined by the corresponding *w*_*i*_. See Figure S3 for a guide on how to interpret *w*_*i*_ as an alignment. D. Multiple sequence alignment between *y*_1_, *y*_2_, *y*_3_ as determined by the variables *w*_1_, *w*_2_, *w*_3_.

**Figure S2:**
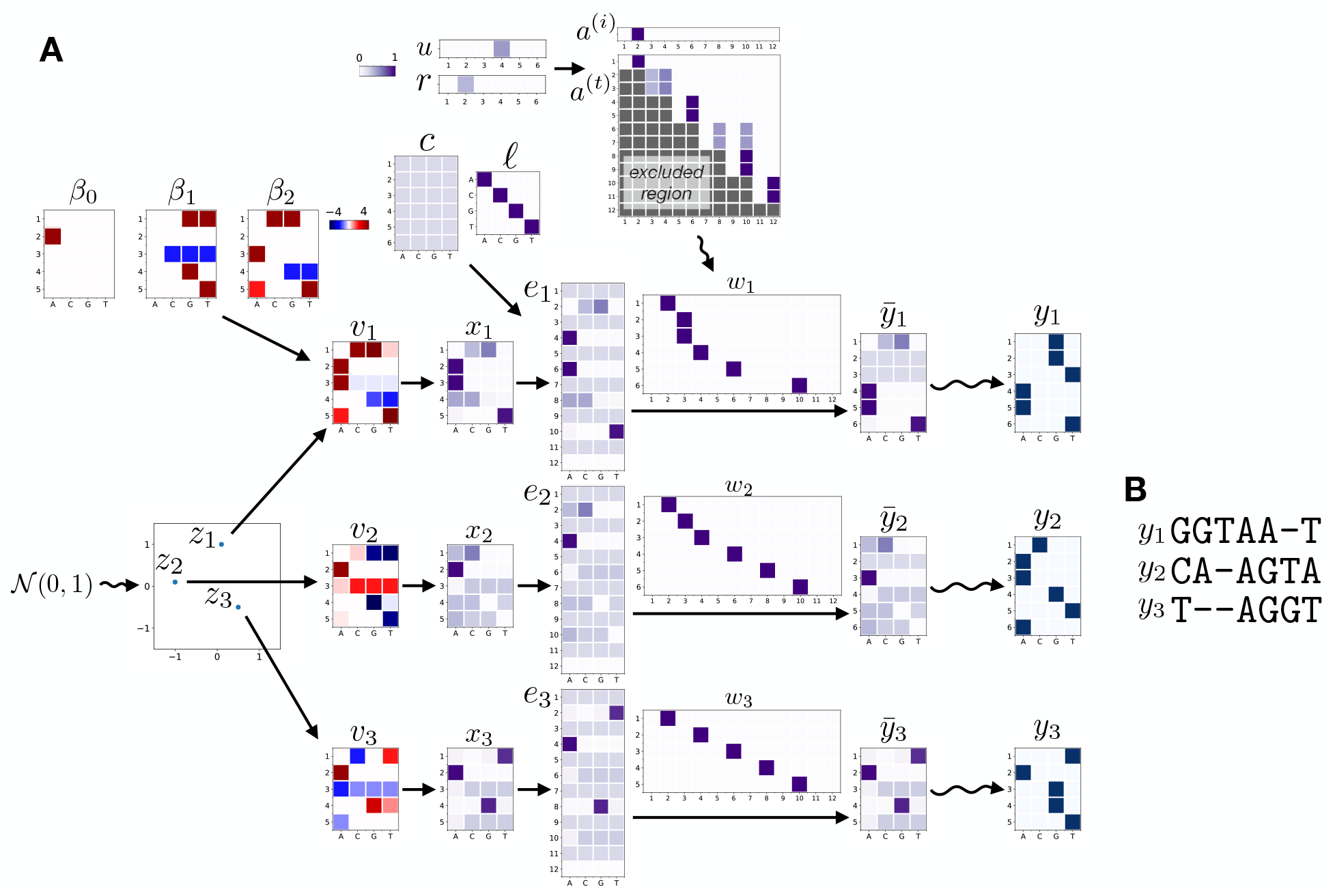
Detailed example of an H-MuE model. A. Illustration of three independent samples from an H-MuE model, with *p*(*v*|*θ*) a probabilistic PCA model with two latent dimensions [1],

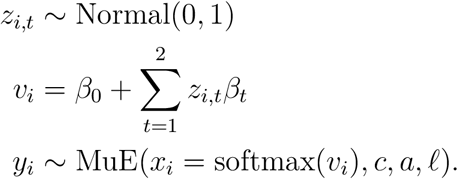

(This is a FactorMuE model with *c* independent of *v*, see Section S5.3.) The diagram components are defined the same way as in Figure S1B. In this example, the generated mutant sequences *y*_1_, *y*_2_ and *y*_3_ are GGTAAT, CAAGTA and `TAGGT`. B. Multiple sequence alignment between *y*_1_, *y*_2_, *y*_3_ as determined by the variables *w*_1_, *w*_2_, *w*_3_.

**Figure S3:**
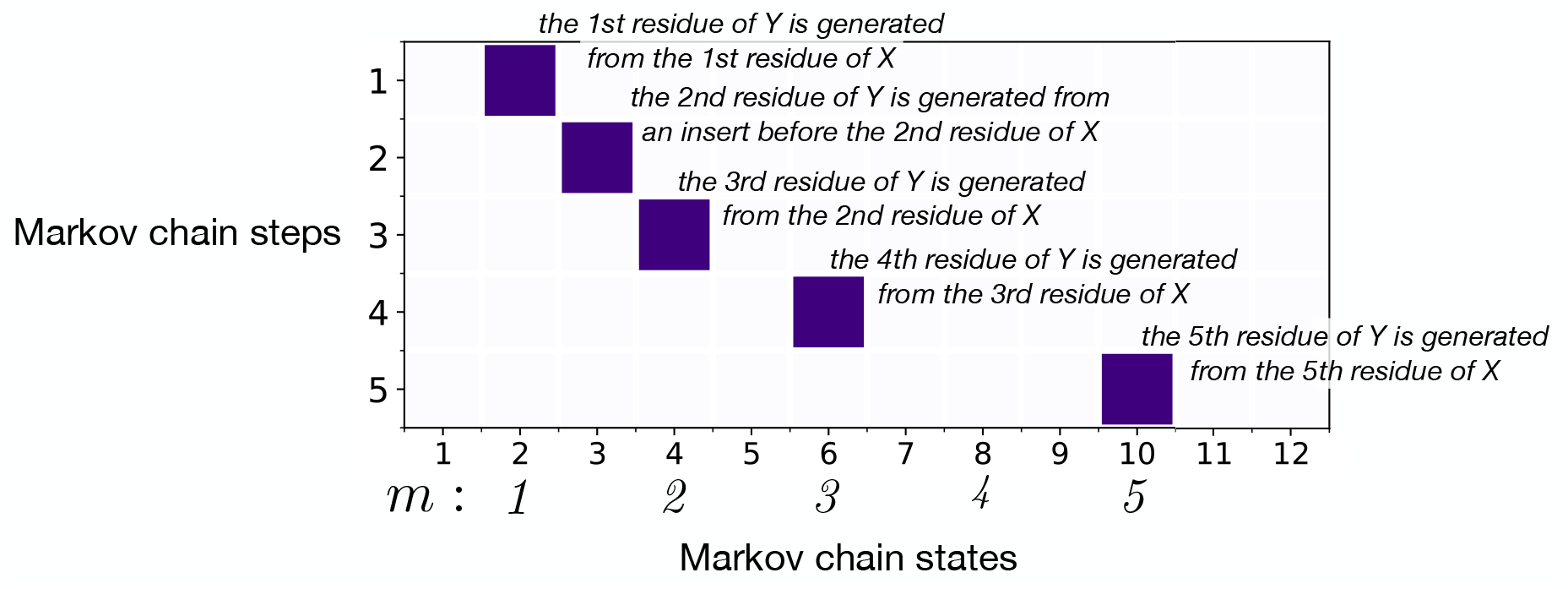
Interpreting the hidden state variable *w*. This plot shows a one-hot encoding of an example *w*; dark purple squares are entries set to 1, and the rest of the matrix is 0. In this example there are *M* = 5 ancestral residues, *K* = 12 states and *L* = 5 residues in *y*. Even-numbered states – that is, match states, with *k*%2 = 0 – are marked below, with their state number *m* = *k/*2. The italic text describes how to interpret each row of the *w* matrix in terms of an alignment. A rigorous mapping from *w* to an alignment is defined in Section S4.1.

**Figure S4:**
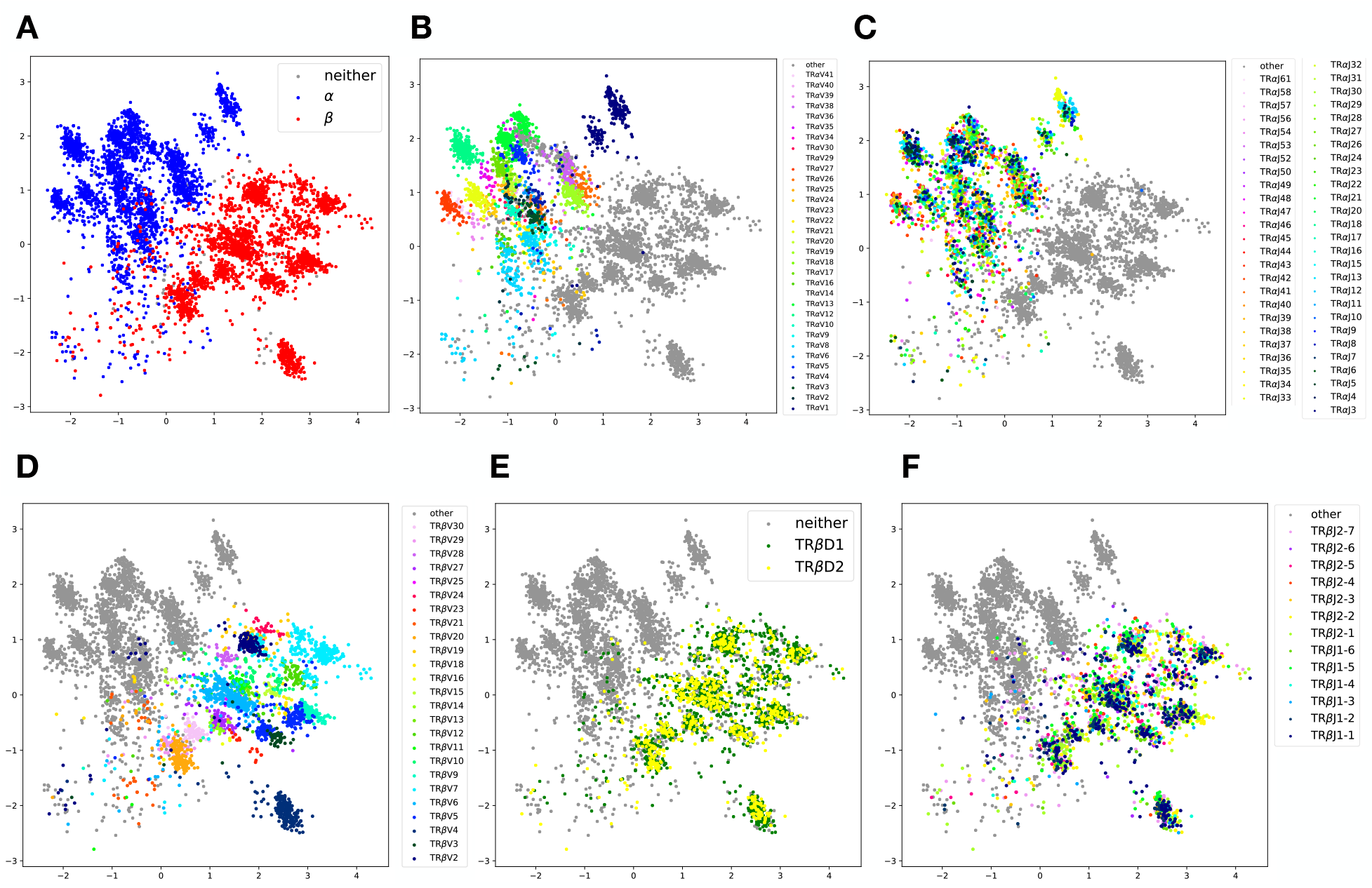
Latent space representation of human T-cell receptor sequences, colored by supervised annotations. Annotations from [35]. A. TCR*α* versus TCR*β*. B. *α* chain V types. C. *α* chain J types. D. *β* chain V types. E. *β* chain D types. F. *β* chain J types and subtypes.

**Figure S5:**
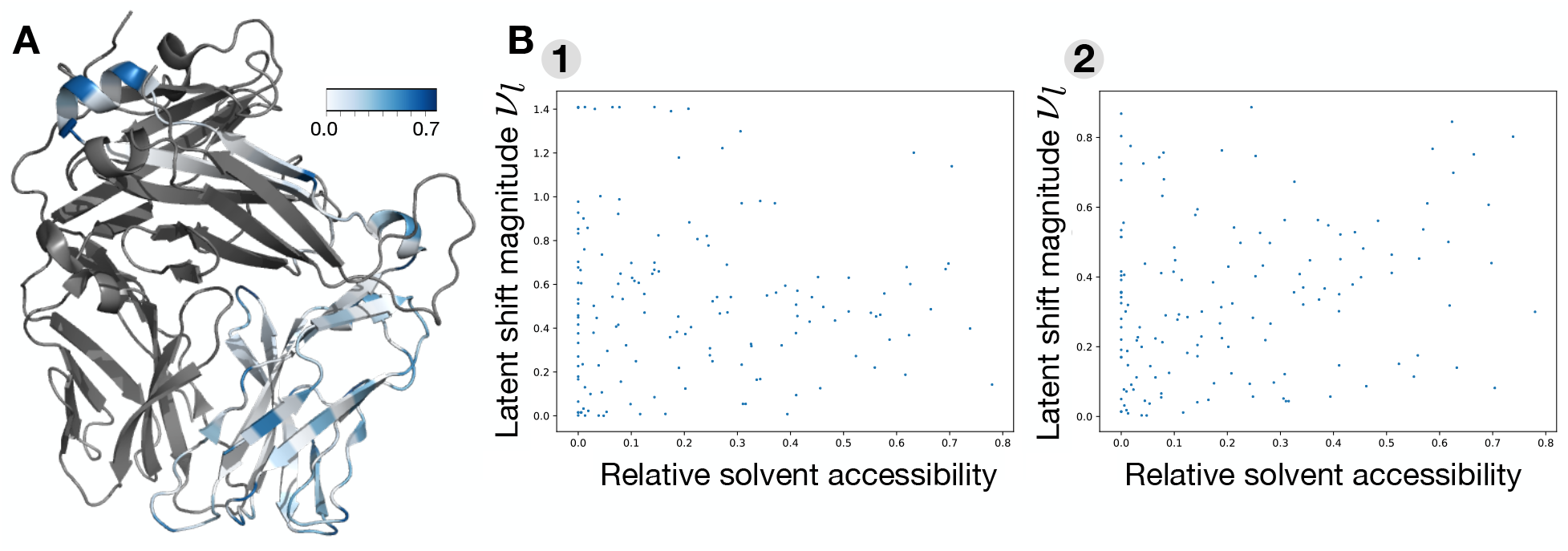
Comparing H-MuE model features to T-cell receptor relative solvent accessibility. A. Relative solvent accessibility of TCR*β* from the structure PDB:2BNR [36] (the TCR*α* chain is shown in grey), computed using DSSP [37] and the maximum values in [38] with the Biopython API [39]. B. Residue relative solvent accessibility versus Factor-MuE shift magnitude *ν*_*l*_ (Equation S78) along vector 1 and vector 2 from Figure 3E. The correlation between the shift along vector 1 and the accessibility is Spearman *ρ* = 0.039, *p* = 0.64.

**Figure S6:**
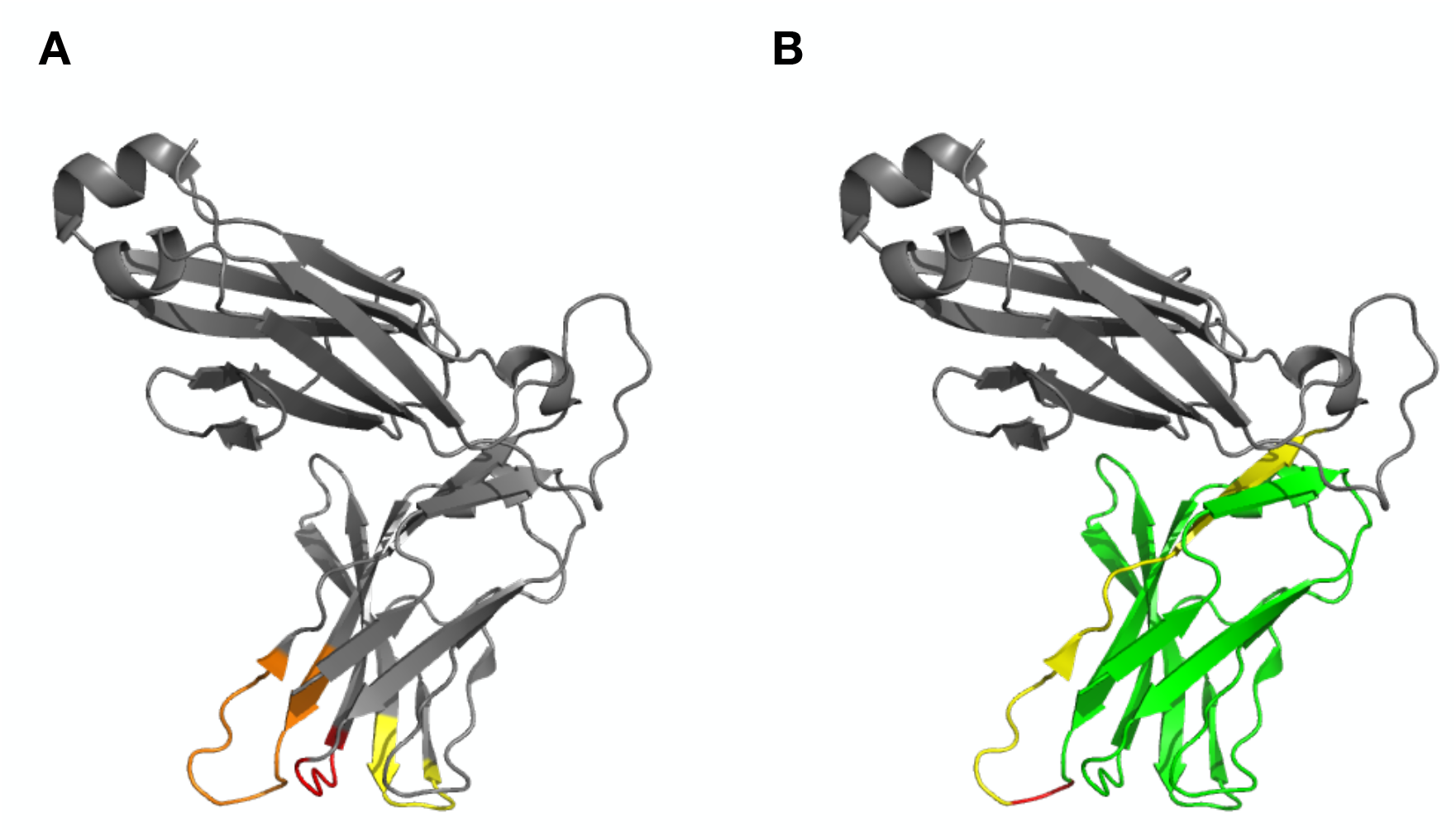
T-cell receptor structural annotations. A. CDR segments of PDB:2BNR chain E [36], based on IgBLAST annotations [40] of the nucleotide sequence of 1G4 TCR*β* obtained from [41], and translated from nucleotides into the corresponding positions in the amino acid sequence. CDR1 in red, CDR2 in yellow and CDR3 in orange. B. V (green), J (yellow) and junction (red) segments of the 1G4 nucleotide sequence, based on the IgBLAST annotations, and translated from nucleotides.

**Figure S7:**
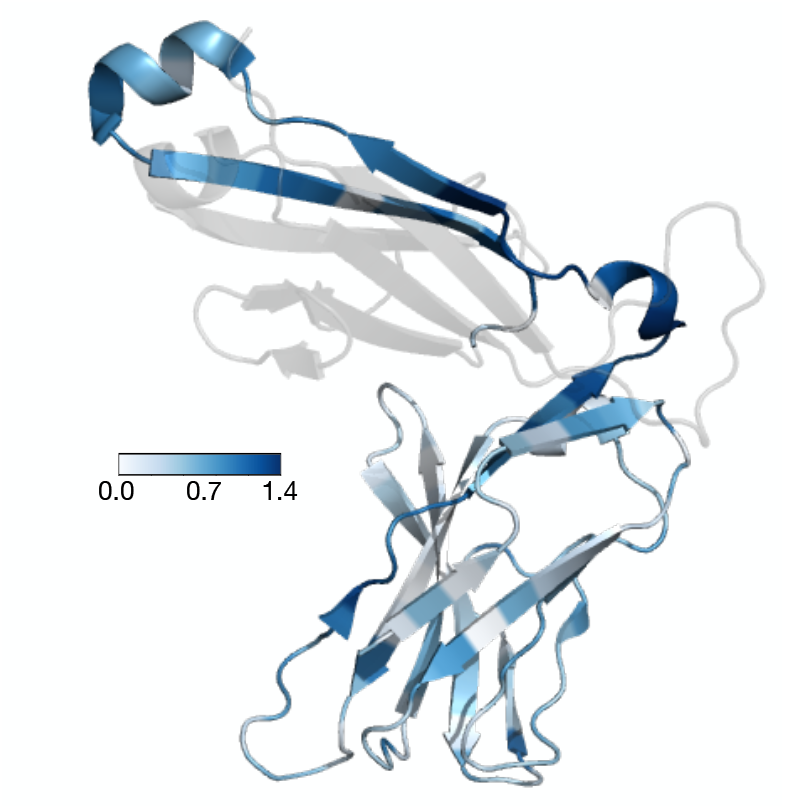
Shift *ν*_*l*_ from chain *α* to chain *β* sequences learned by the RegressMuE model. *ν*_*l*_ was computed as in Equation S78, using the chain index in place of the latent variable *z*.

**Figure S8:**
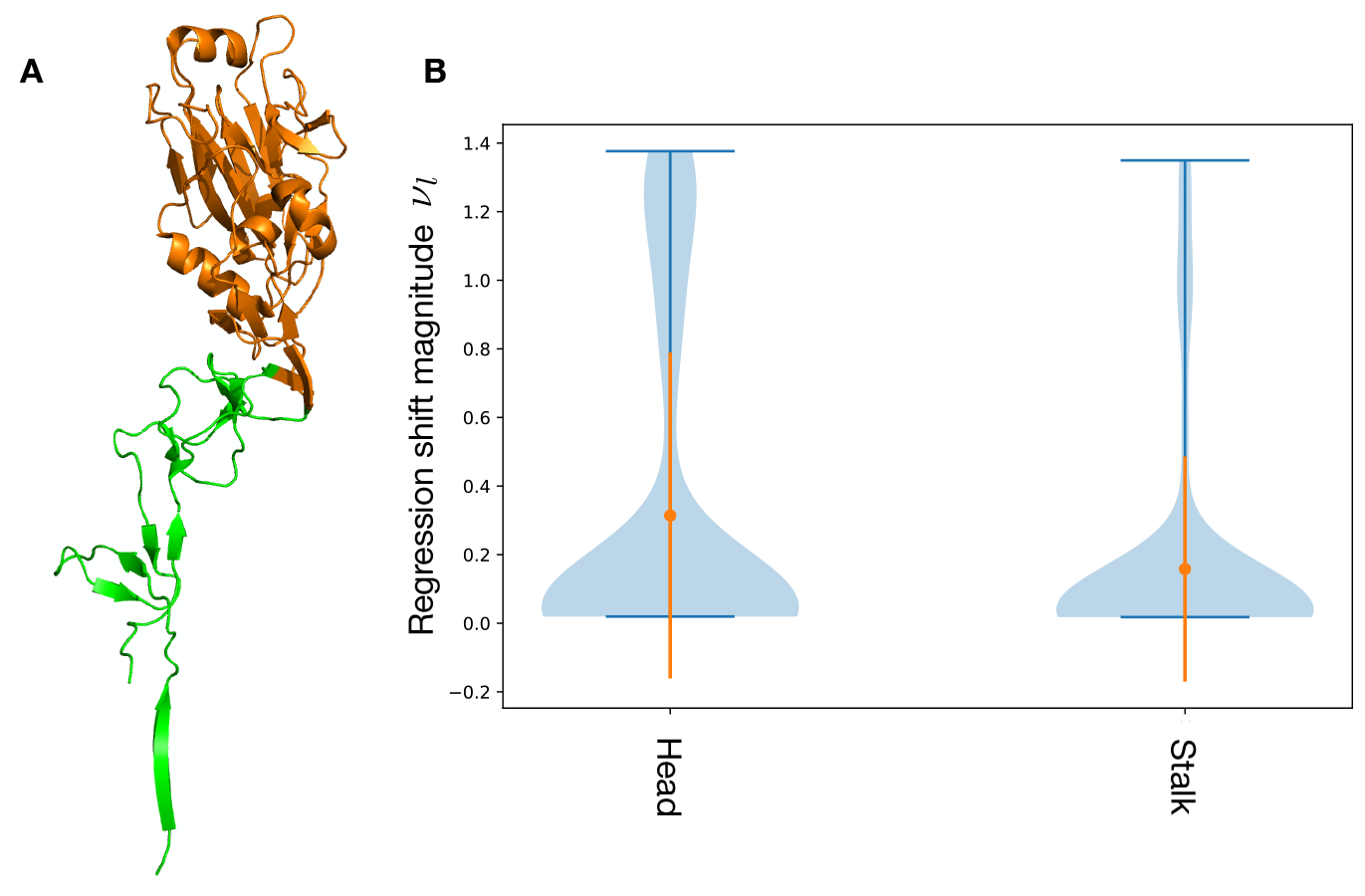
Comparing H-MuE model regression coefficients to HA1 structural domains. A. Head (orange) and stalk (green) domains of the HA1 protein (PDB:4O5N); residues between sites 52 and 277 are defined as the head domain, and all others as stalk, following [42]. B. Violin plots of regression shift *ν*_*l*_ (Equation S80) for residues in the head domain (226 residues) versus the stalk domain (103 residues). Mean and standard deviation are shown in orange.

**Figure S9:**
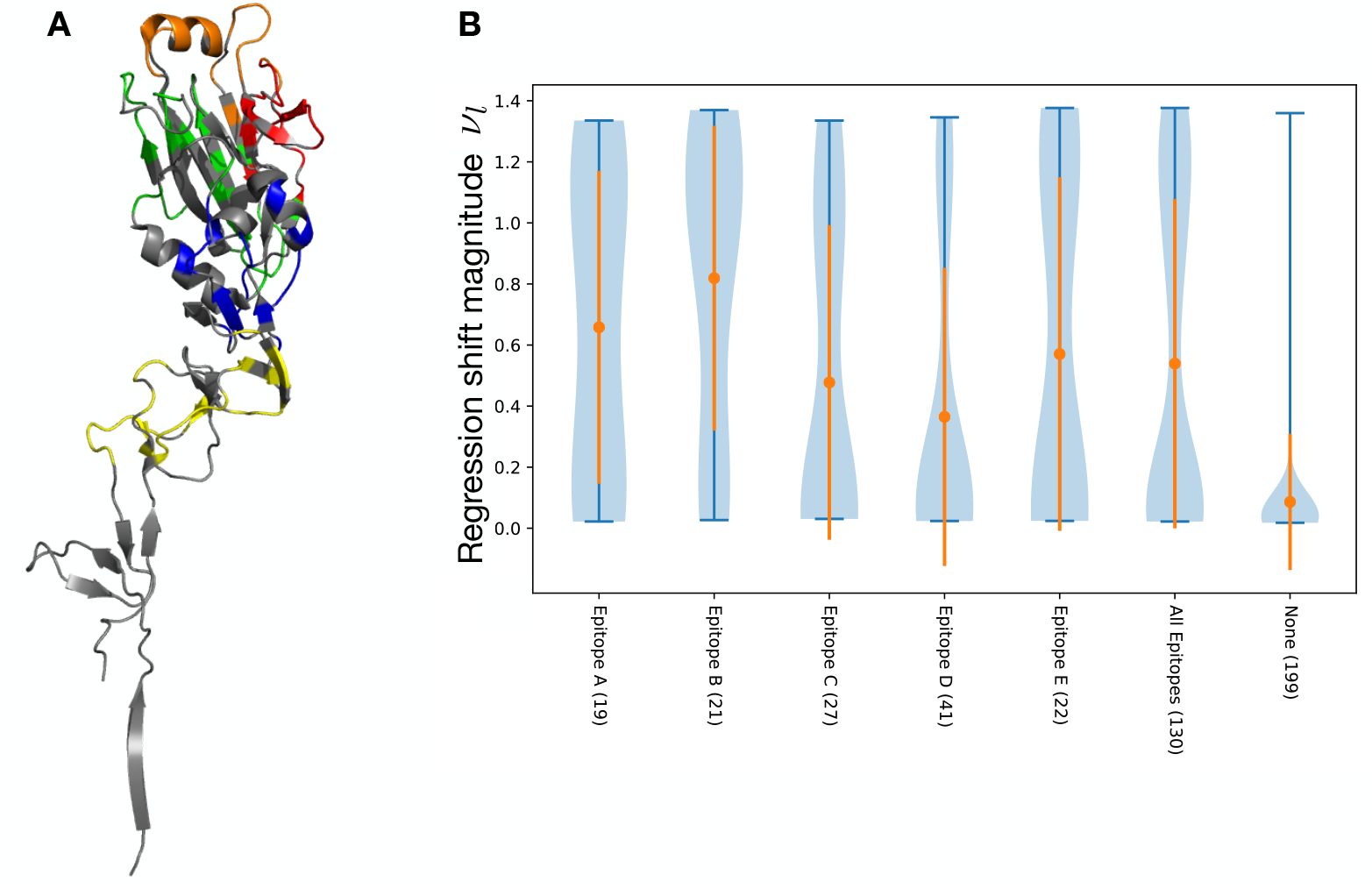
Comparing H-MuE model regression coefficients to HA1 epitope regions. A. Epitope regions A (red), B (orange), C (yellow), D (green), E (blue) [25, 27]. B. Violin plots of regression shift *ν*_*l*_ (Equation S80) for residues in each epitope region, for all epitope regions together, and for residues not in any epitope region; the number of residues in each region is shown in parenthesis. Mean and standard deviation are shown in orange.

**Figure S10:**
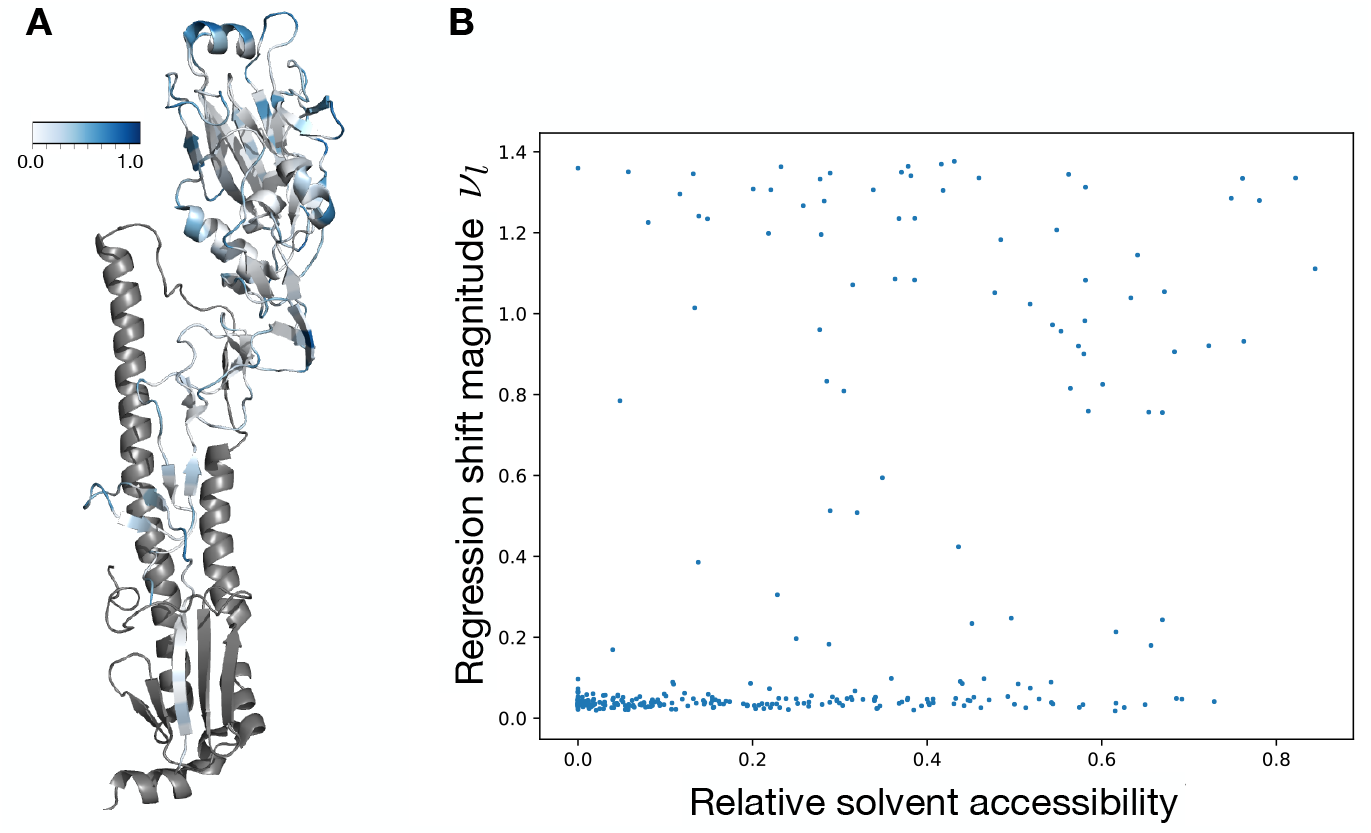
Comparing H-MuE model regression coefficients to HA1 relative solvent accessibility. A. Relative solvent accessibility of the HA1 protein (PDB:4O5N), computed using DSSP [37] and the maximum values in [38] with the Biopython API [39]. HA2 protein shown in grey. B. Relative solvent accessibility versus regression shift magnitude *ν*_*l*_ (Equation S80), residue-by-residue. Spearman *ρ* = 0.41, *p* < 1*e* − 13.

**Figure S11:**
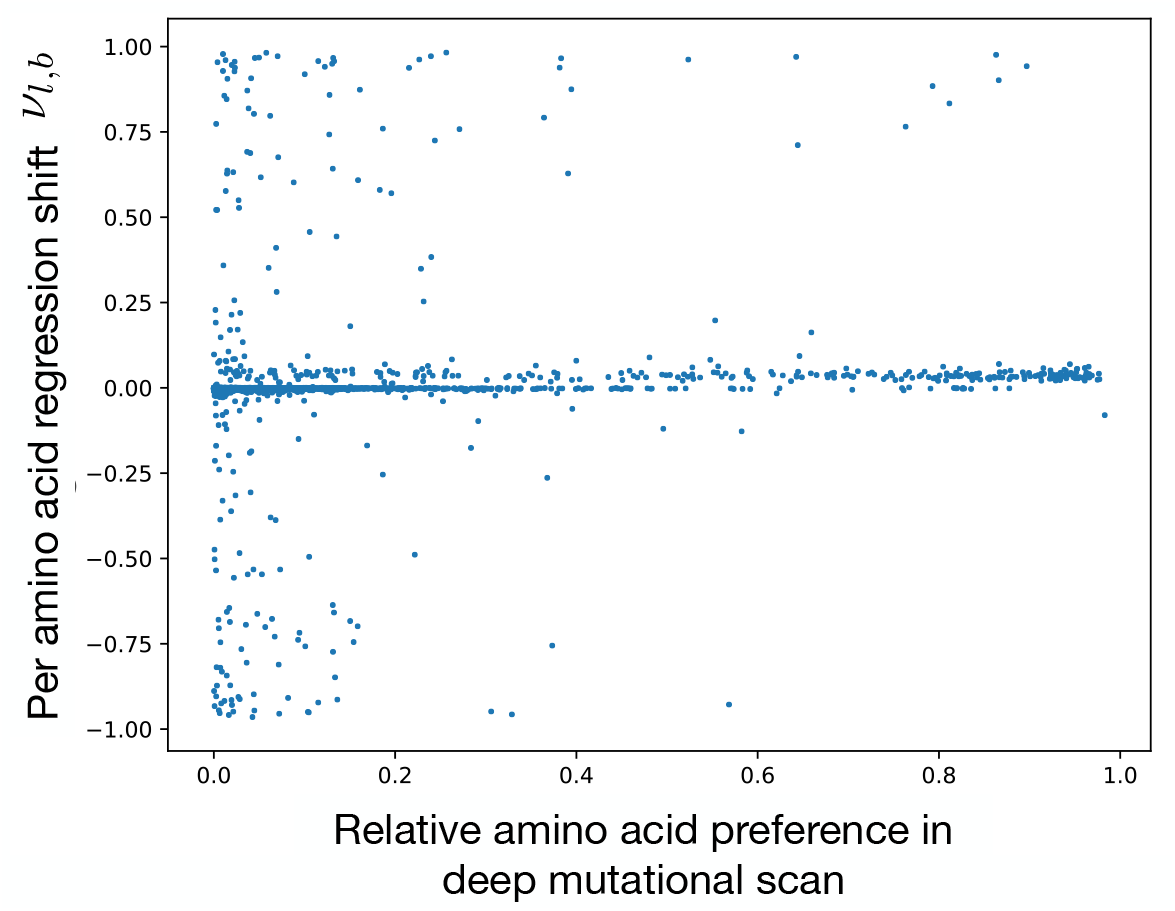
Comparing H-MuE model regression coefficients to a deep mutational scan of HA. X-axis: regression shift for each amino acid at each position from 1968 to 2019,

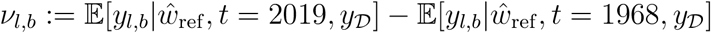

(terms defined as in Equation S80). Y-axis: relative preference for point mutants with amino acid *b* at position *l* in the deep mutational scan performed in Lee et al. [42]. Spearman *ρ* = 0.08, *p* < 1*e* − 11.

### S2 Statistical Pathologies in MSA-based Methods

In this section we describe the theoretical problems that arise when using MSA-based methods for predicting biological sequences. The underlying issue is that the multiple sequence alignment of the entire dataset may change as more data is added, since new sequences may contain regions that do not match old sequences, and since new sequences provide evidence for and against similarity among old sequences. The fact that the shape of the data matrix may change as more data is added and the fact that previously observed data may be altered as more data is added make random variables and predictions ill-defined.

More formally, let 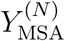 be the alignment of a dataset of *N* sequences, with 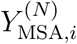 corresponding to the alignment of the *i*th sequence. Let 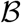 be the symbol alphabet including the gap symbol, e.g. for DNA 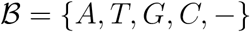. Let *J*^(*N*)^ be the width of the alignment. Each row of the alignment is a vector of symbols 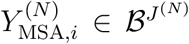. The standard method of building probabilistic models of pre-aligned sequence data is to assume that aligned sequences 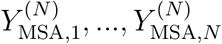 are independently and identically distributed (iid) according to some underlying distribution 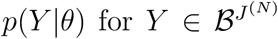, such as a Potts model or neural network [14, 17, 43]. Mathematically,

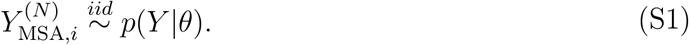

While models of this form are powerful when used for parameter estimation (that is, inferring *θ*), they are ill-defined when used for prediction of unobserved or future sequences.

The posterior predictive distribution 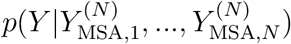 is defined for 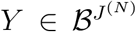. However, when we observe a new sequence and add it to our multiple sequence alignment, we have in general *J*^(*N*+1)^ ≠ *J*^(*N*)^ and consequently 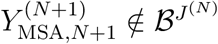. Thus we find that

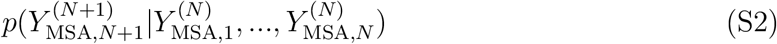

is ill-defined. Future datapoints will not necessarily live in the same mathematical space as the model.

Even if we could somehow safely assume that *J*^(*N*+1)^ = *J*^(*N*)^ we would still run into problems, since the alignment can change with additional data. Mathematically, in general 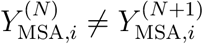 for *i* ∈ {1, …, *N*}. Then,

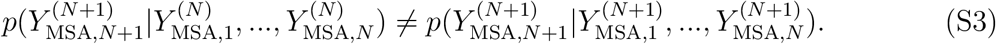

In other words, making predictions of unobserved data (the left hand side of the equation), is not the same as holding out and predicting pre-aligned data (the right hand side of the equation).

Ultimately, both of these problems emerge because pre-aligned sequence data automatically violate the iid assumption of the model in Equation S1. The aligned sequence 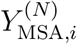 is not just a function of the unaligned sequence *y*_*i*_, it is also a function of all the other unaligned sequences *y*_*i*′≠*i*_ in the dataset. In theory, we could avoid prediction problems by specifying a joint model of the entire dataset, treating the entire MSA as a stochastic object that changes with an increasing number of sequences *N*. That is, we could define a model 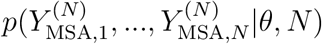, where in general 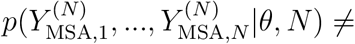 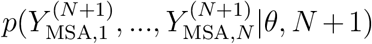. While such models are possible in theory, in practice they are challenging to write down, hard to perform inference on, and lack the asymptotic convergence guarantees associated with iid models. H-MuE models offer a principled alternative that preserves the iid assumption while making sequence prediction well-defined.

Note also that prediction is a core concern across all of statistics, not just in forecasting applications. Many model evaluation methods, such as cross validation and posterior predictive checks, depend on the ability of statistical models to predict unobserved data [44]. Prequential statistical approaches depend entirely on prediction [45]. Causal inference methods are founded in prediction [46]. In general, relying on multiple sequence alignments makes application of these powerful ideas ill-defined. H-MuE models make application possible.

### S3 MuE Distribution Details

#### S3.1 Termination

The state variable *w* in the MuE distribution is drawn from a Markov chain. We consider two possible methods for terminating the chain. The first method is to have an explicit termination state: when the Markov chain reaches the final state *k* = *K*, it halts without emitting an observation. This method is used in the previously proposed probabilistic alignment models discussed in Section S4. The second method is to sample the length of the chain independently of the states of the chain. We use this approach in our H-MuE models, since it makes inferring an accurate value for *M* unnecessary, and we can instead simply choose a large *M* value compared to the typical length of sequences in the dataset (Section S8).

#### S3.2 Special Cases

In this section we describe two important special cases of the MuE distribution. First, consider the case where 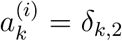 and 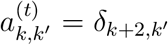, where δ is the Kronecker delta. If, in addition, *L* = *M* (which will be guaranteed if we are working with an explicit termination state *k* = *K*), then the MuE distribution simplifies to

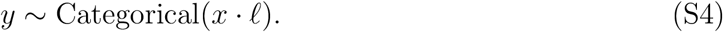

We refer to this as the “no-indel case” of the MuE distribution: the generated *y* sequences just have substitution mutations relative to *x*.

If, in addition, we set *D* = *B* and *ℓ* = *I*_*B*_, where *I*_*B*_ is the *B* × *B* identity matrix, then *y* ∼ Categorical(*x*) under the MuE distribution, and the full H-MuE model becomes

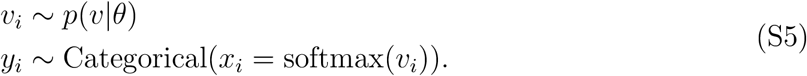

We thus recover the standard multivariate categorical emission distribution, used in e.g. multinomial logit regression models [47] and the state-of-the-art pre-aligned sequence model proposed by Riesselman et al. [14]. We refer to this as the “no-mutation case” of the MuE distribution.

### S4 Theory

In this section we describe our theoretical results. First we rigorously establish a mapping between the latent variable *w* and a pairwise alignment between *x* and *y*, and show that the restrictions on the transition matrix *a*^(*t*)^ are necessary and sufficient for this mapping to exist with probability one (Section S4.1). We then show that the MuE distribution provides a unified framework for understanding a wide variety of classical biological sequence analysis methods: we can recover, as special case models and estimators, stochastic process models of sequence evolution (the Thorne-Kishino-Felsenstein model, Section S4.2), probabilistic multiple sequence alignment methods (the profile HMM, Section S4.4), probabilistic pairwise sequence alignment methods (the pair HMM, Section S4.3), and non-probabilistic pairwise sequence alignment algorithms (the Needleman-Wunsch algorithm, Section S4.5). We also compare the MuE distribution to a natural language translation model (Section S4.6), which fits the basic form of the MuE distribution but violates the restrictions on *a*^(*t*)^.

#### S4.1 Alignments and States

For convenience, we assume throughout Section S4.1 that the Markov chain terminates at the state *k* = *K* = 2*M* + 2, and that it reaches the termination state eventually with probability one. A precisely analogous analysis can be done for the case where there is no termination state and instead *L*, the length of sequence *y*, is drawn independently of the Markov chain.

##### S4.1.1 Definitions

In this section we set up key definitions, illustrated with the example in Equation S6, S7 and S8.

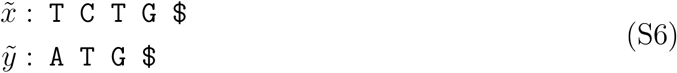

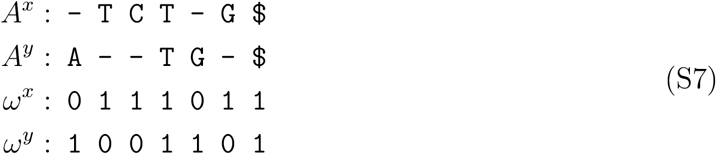

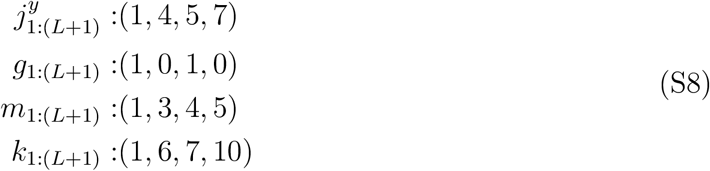

Let 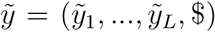 and 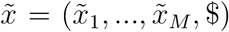 be sequences, with $ the termination symbol. A pairwise alignment 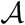 of 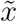 and 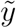 with *J* columns is defined by the tuple (*A^x^, A^y^*) where *A*^*x*^ is a vector of length *J* consisting of the residues of 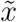, in order, interspersed with gap symbols, and similarly for *A*^*y*^, with the condition that every column of 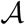 contains either a letter from 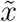, a letter from 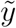, or both, and the final column must contain the termination symbol from both sequences. We can represent the alignment in terms of the index vectors ω^*x*^, ω^*y*^ ∈ {0, 1}^*J*^, where 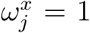 if there is a letter of 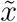 at 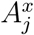 and 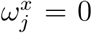 if there is a gap, and likewise for ω^*y*^ and *A*^*y*^. If

1. 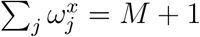. (The sequence 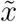 has *M* residues plus the termination symbol.)
2. 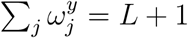. (The sequence 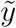 has *L* residues plus the termination symbol.)
3. 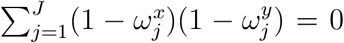. (Each column of the alignment has at least one residue; there cannot be two gap symbols aligned.)
4. 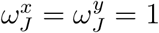. (The final column contains the terminal symbol of each sequence.)

then the tuple (ω^*x*^, ω^*y*^) uniquely defines an alignment of the sequences 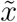 and 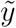. Let 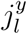, for *l* ∈ {1, …, *L* + 1}, index the column of the alignment that the *l*th residue of 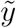 falls in. Mathematically,

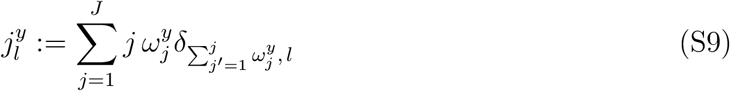

where δ is the Kronecker delta. Let *g*_*l*_ indicate whether the *l*th residue of 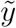 is aligned to a gap in 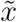 or not. Mathematically,

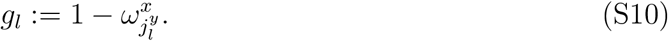

Let *m*_*l*_ be the residue of 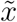 aligned to the *l*th residue of 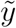; if the *l*th residue of 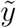 is aligned to a gap in 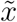, let *m*_*l*_ be the closest residue in 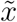 to the right. Mathematically,

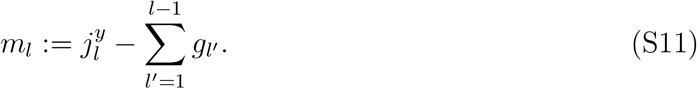

Finally, define

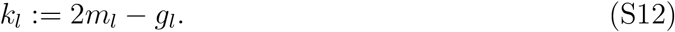

##### S4.1.2 From Alignments to States

Starting from any pairwise alignment 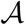 we can compute *k*_*l*_ for *l* ∈ {1, …, *L* + 1}. We can use these *k*_*l*_ to define a sequence of states *w* for the Markov chain in a MuE distribution, by setting 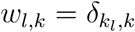.

Alignments are typically intended to represent an evolutionary relationship between sequence 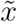 and 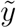. If residue *l* of 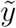 is aligned to residue *m* of 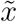, it suggests that they share common descent, and should therefore be similar. In particular, we can expect 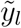 to either match 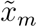 exactly or be related via a substitution mutation. The states *w* defined from the alignment 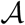 reflect this idea. If the *l*th residue of 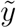 is aligned to the *m*th residue of 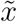, then *m*_*l*_ = *m* and 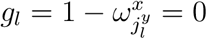. The MuE distribution gives

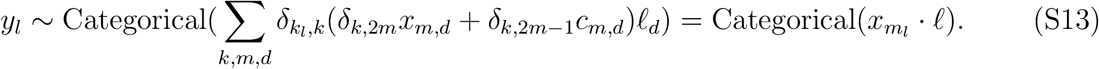

Thus, *y*_*l*_ is generated from 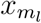 according to the substitution probabilities *ℓ*. On the other hand, if the *l*th residue of 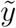 is aligned to a gap, 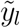_*l*_ should be independent of 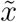, since it comes from an insertion mutation. In this case 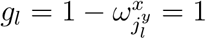 and we find

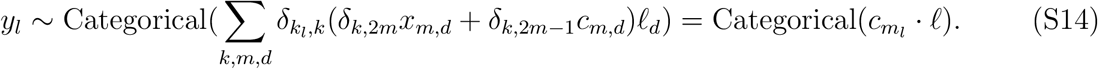

Thus, *y*_*l*_ does not depend on *x* and is instead determined by the insertion parameter *c*.

##### S4.1.3 From States to Alignments

Starting from a sequence of states *w* drawn from MarkovModel(*a*^(*i*)^, *a*^(*t*)^), we can convert uniquely back to an alignment 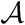 between 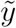 and 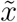. We assume *w*_*L*+1_ is in the termination state, ie. *w*_*L*+1,*K*_ = 1 (note that this assumption requires that the Markov chain reaches the termination state with probability one). Let *k*_*l*_ = arg max*_k_ w_l,k_* be the state of the Markov model at position *l*. We can now invert Equations S9-S12 to reconstruct the corresponding alignment. We have

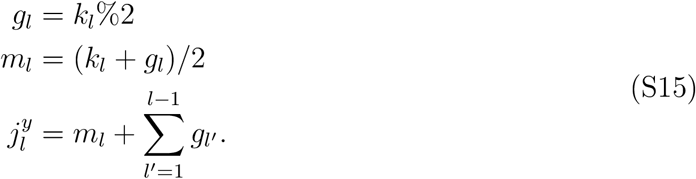

where recall % is the modulo operation. (Note that for *j*^*y*^, the list of columns of the alignment that the residues of 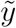 fall in, to be valid based on its definition, it must be a strictly ascending list of integers; this is discussed further below.) We then obtain, for *l* ∈ {1, …, *L* + 1},

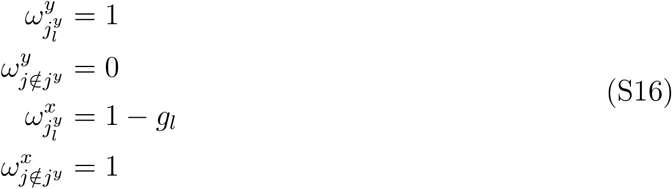

where we have used the property that 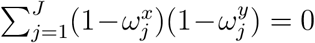. ω^*x*^ and ω^*y*^ together uniquely define an alignment 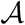 of 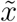 and 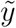.

As mentioned above, for *j*^*y*^ to be valid based on its definition, it must be a strictly ascending list of integers. That is, we must have for all *l* ∈ 2, …, *L* + 1,

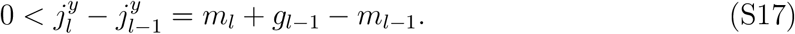

Intuitively, this condition says that alignments cannot “double back”: if residue *m* in 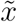 is aligned to residue *l* in 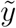, then a later residue *l′ > l* cannot align to an earlier residue *m′ < m* in *y*. With some more algebra, we find that for all *l* ∈ 2, …, *L* + 1 we must have

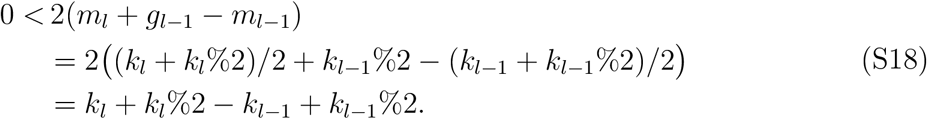

For this condition to hold with probability one for any sample from the Markov chain it must be the case that the transition probability 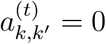 for *k′* + *k′*%2 − *k* + *k*%2 ≤ 0, so long as state *k* is accessible by the Markov model (that is, so long as the Markov chain can eventually reach state *k*, starting from the initial state). The MuE distribution has this restriction, and therefore every sample from a MuE distribution corresponds to an alignment.

##### S4.1.4 Multiple Sequence Alignments

When we have multiple independent samples *y*_1_, …, *y*_*N*_ from a MuE distribution or H-MuE model, the collection of associated state variables *w*_1_, …, *w*_*N*_ can be interpreted as a multiple sequence alignment of *y*_1_, …, *y*_*N*_. While there is some ambiguity here in the mapping between states and alignments (unlike in the pairwise alignment case), we present the mapping used with the profile HMM, relying on the re-derivation of the pHMM as a special case of the MuE distribution described in Section S4.4. The core idea is to interpret the positions of *x* as conserved columns. Let *k_i,l_* = arg max *w_i,l,k_* be the state the Markov chain is in at position *l* in sequence *i*, with *m_i,l_* and *g_i,l_* defined following Equation S15. For each *l*, if *g_i,l_* = 0, we place *y_i,l_* in the *m_i,l_*th conserved column of the MSA. Otherwise, if *g_i,l_* = 1, we place *y_i,l_* into the block of insertions immediately before the conserved column *m_i,l_*. As described in detail in Durbin et al. [16] Chapter 6.5, the choice of how to represent the insertion blocks in the alignment is somewhat arbitrary. The standard approach is to pad the insertion block on the right with gap symbols to reach the length of the longest subsequence in state *k_i,l_* across the entire dataset. See Figure S2 for an example and Durbin et al. [16] Chapter 6.5 for further details.

#### S4.2 Thorne-Kishino-Felsenstein

The Thorne-Kishino-Felsenstein (TKF) model is a continuous-time stochastic process model of sequence evolution [15]. Let *x* be a one-hot encoding of the initial sequence (excluding the termination symbol). Let *D* = *B* and let π be the TKF parameter corresponding to the equilibrium probability of each letter. For all *m* ∈ {1, …, *M* + 1} and *b* ∈ {1, …, *B*}, assign

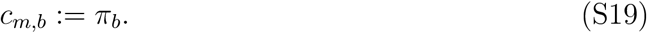

Let λ > 0 and *μ* > 0 be the TKF indel rate parameters, with λ *< μ*, and let *τ* > 0 be the divergence time parameter. Define

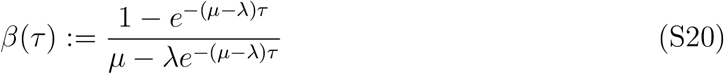

It is convenient to index states *k* and *k′* using the corresponding (*m, g*) values (Equation S15), ie. *m* = (*k* + *k*%2)/2, *g* = *k*%2, *m′* = (*k′* + *k′*%2)/2 and *g′* = *k′*%2. Then we define the transition matrix

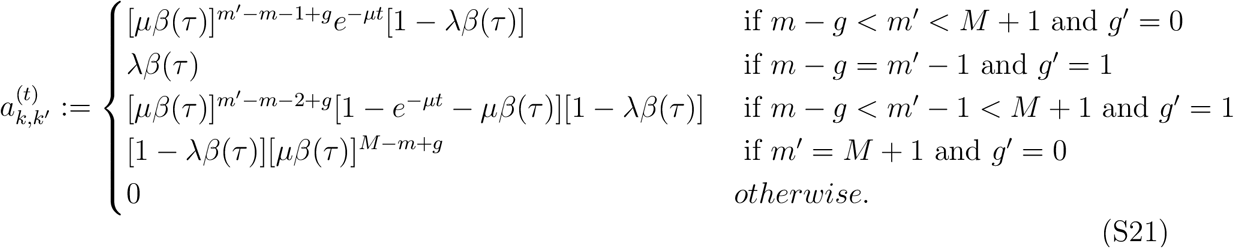

The initial transition vector follows the same form, and can be written as 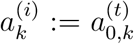. The termination state is the final state *k* = *K* = 2*M* + 2. Let *s* > 0 be the TKF substitution rate parameter and define the substitution matrix

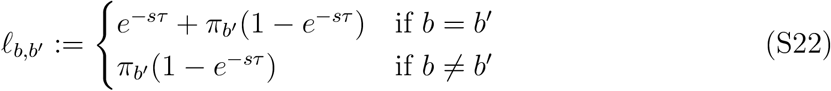

With these definitions, *y* ~ MuE(*x, c, a*, *ℓ*) is the transition distribution of the Thorne-Kishino-Felsenstein model after the sequence *x* evolves for time *τ*.

Note that in the limit λ*,μ* → 0 we recover the no-indel special case of the MuE (Section S3.2). In the limit *τ* → 0 we recover the no-mutation case, and *y* = *x* with probability one. Figure S12 illustrates samples from the TKF model with changing parameters.

**Proof**

We will show that the joint probability of *w* and *y* under the MuE distribution is identical to the joint probability of the corresponding alignment and *y* under the TKF model. To start, we systematically enumerate state transitions in the MuE model and compute the corresponding probability factor under the Thorne-Kishino-Felsenstein (TKF) alignment scoring system. Our alignment notation in this section follows Thorne et al. [15]. “X” represents a residue and “-” a gap. (This notation is equivalent to the ω^*x*^, ω^*y*^ pairwise alignment notation from Section S4.1, with X corresponding to a 1 in ω and - corresponding to a 0.) “.” represents the “immortal link” in the model, the start of the sequence. We use “$” as a termination symbol. Following the original paper, we define, for *ν* ∈ {1, 2, *…*},

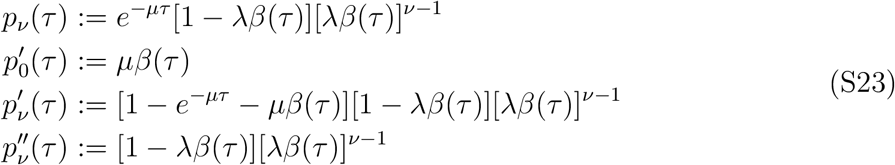

**Figure S12:**
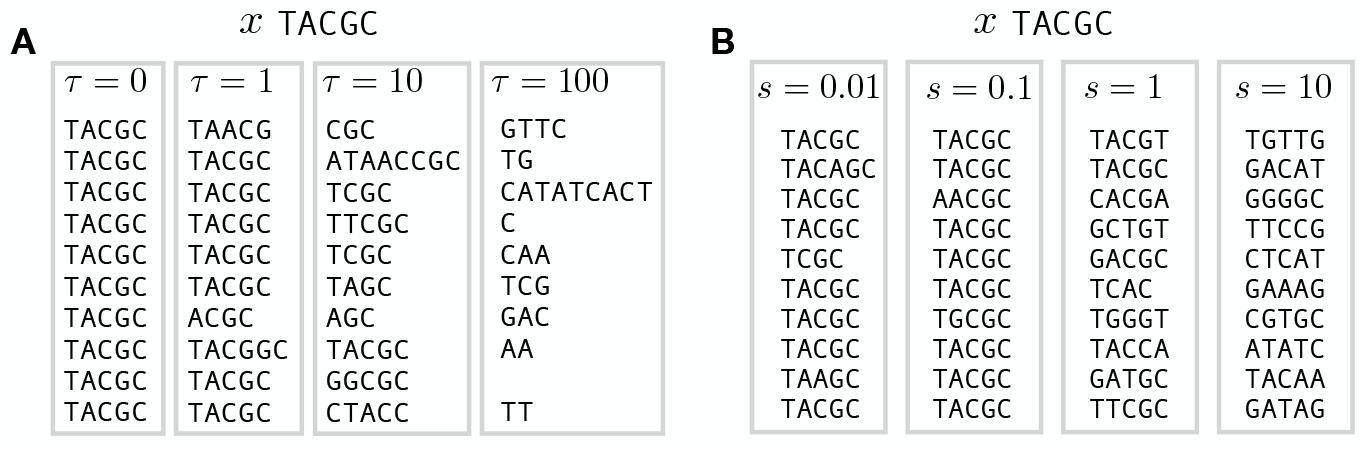
Samples from the Thorne-Kishino-Felsenstein model. Initial sequence `TACGC`, *μ* = 0.02, and λ = 0.01. A. *s* = 0.01 and varying *τ*. B. *τ* = 1 and varying *s*.

As before, we refer to states *k* by tuples (*m, g*), where *m* = (*k* + *k*%2)/2 and *g* = *k*%2. The TKF model assigns probabilities to a pairwise alignment based on the pattern of residues and gaps; we enumerate all possible state transitions in the MuE Markov model and compute the probability factor that they contribute under the TKF scoring system. When enumerating transitions in the Markov model we put a “|” symbol to the right of the residue we are transitioning *from*.

1. Transitioning from a state (*m*, 0) to a state (*m′ > m*, 0) gives the probability factor 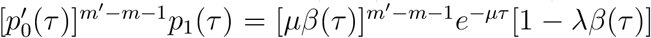 according to the TKF scoring system.

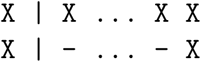
2. Transitioning from (*m*, 1) to (*m′ ≥ m*, 0) gives the factor 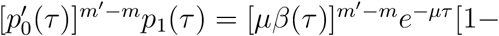 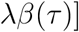.

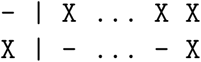
3. Transitioning from (*m*, 1) to (*m*, 1), situation 1. This gives a factor 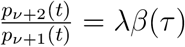.

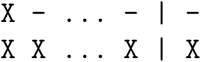
4. Transitioning from (*m*, 1) to (*m*, 1), situation 2. This gives a factor 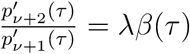

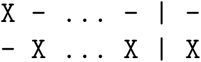
5. Transitioning from (*m*, 0) to (*m* + 1, 1). This gives a factor 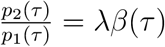.

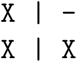
6. Transitioning from (*m*, 0) to (*m′ > m* + 1, 1). This gives a factor 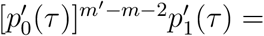 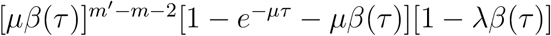.

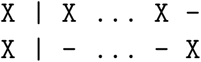
7. Transitioning from (*m*, 1) to (*m′ > m*, 1). This gives a factor 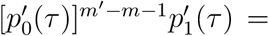 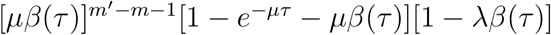.

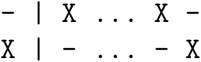
8. Transitioning from (*m*, 0) to (*M* + 1, 0). This gives a factor 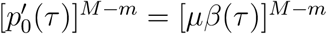.

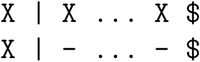
9. Transitioning from (*m*, 1) to (*M* +1, 0). This gives a factor 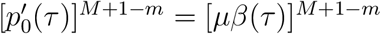.

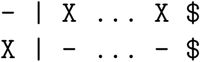
10. Initial transition to (1, 1). This gives a factor 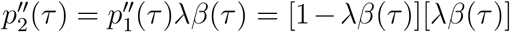.

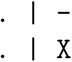
11. Initial transition to (*m*, 0). This gives a factor 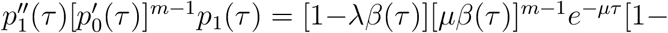 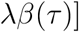.

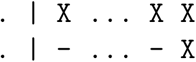
12. Initial transition to (*m* > 1, 1). This gives a factor 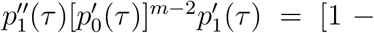 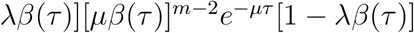.

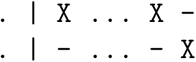
13. Initial transition to (*M* + 1, 0). This gives a factor 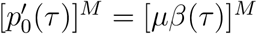.

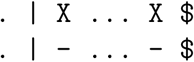

Compiling these results yields the probability factors associated with each transition between states

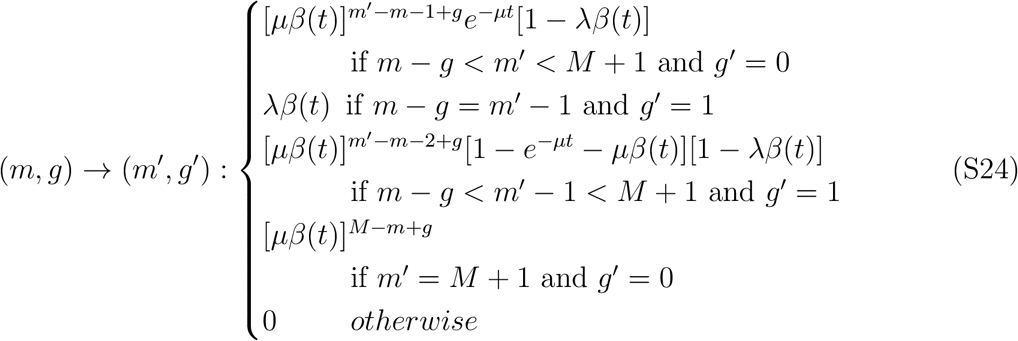

And with each initial transition

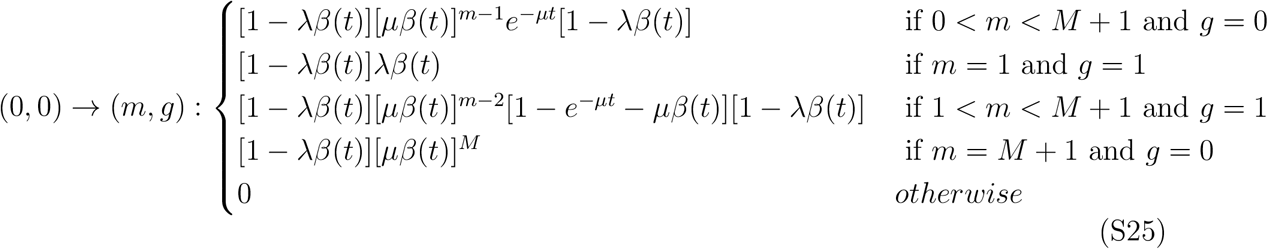

However, these are unnormalized probability factors, not complete probabilities. Note that every alignment will have a factor (1 − λ*β*(*t*)), which in the original TKF description is associated with the initial transition. However, if we instead rearrange this factor and assign it to the final transition we obtain the transition matrix given in Equation S21. We can check that this transition matrix normalized. From a state (*m*, 0), the total outward transition probability is one:

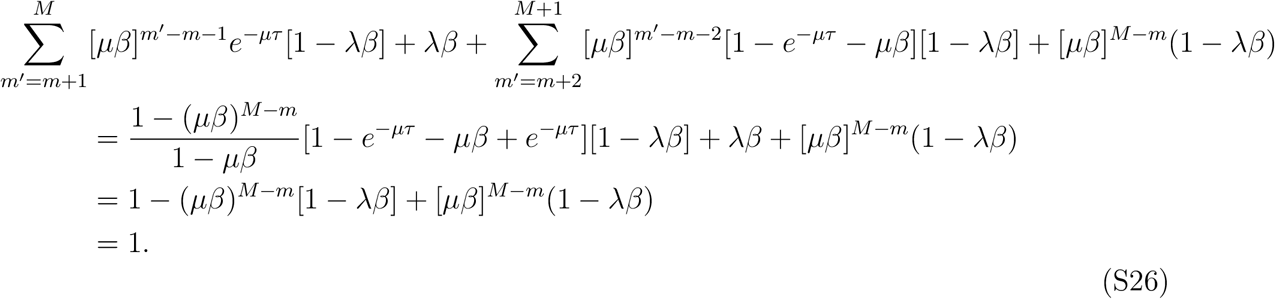

The same expression holds for the initial transition, plugging in *m* = 0. From (*m*, 1), we have

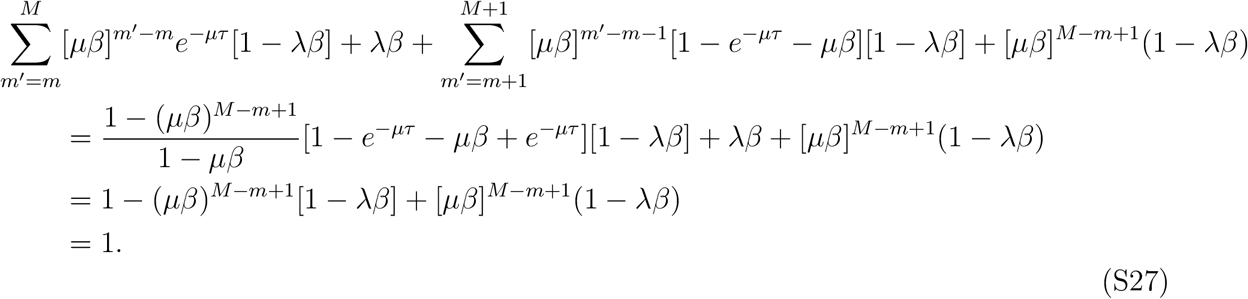

Conditional on the *m*th residue of *x* being aligned to the *l*th residue of *y* (i.e. *w_l,_*_2*m*_ = 1), the TKF model specifies that the probability of *y*_*l*_ given *x*_*m*_ is ∑_*b,b′*_*x*_*m,b*_*ℓ*_*b,b′*_ *y*_*l,b*′_, which is identical to the probability under the MuE model. In the case where the *l*th residue of *y* is aligned to a gap (ie. *w_l,_*_2*m*−1_ = 1), the TKF model says the probability of choosing the specific base *b* is π_*b*_, the equilibrium probability of the base. We can check that the MuE provides the same factor:

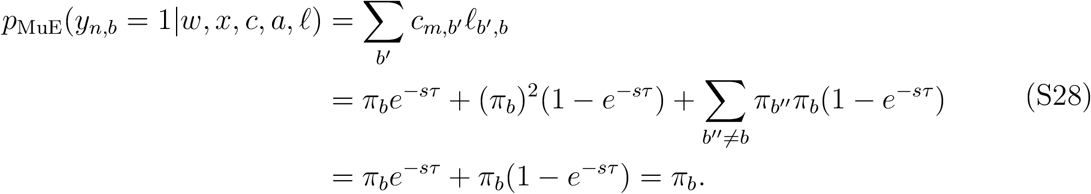

#### S4.3 Pair HMM

The pair HMM model generates pairwise alignments by switching between three states: (1) a state emitting residues in both *x* and *y* (a match state), (2) a state emitting a residue in *x* and a gap in the alignment of *y*, and (3) a state emitting a gap in the alignment of *x* and a residue in *y*. Figure S13 shows a standard pair HMM diagram and state probabilities, with γ the probability of transitioning to a gap state, *ϵ* the probability of staying in a gap state, and *κ* the probability of the Markov chain terminating [16]. We assume 1 − 2γ − *κ* ≥ 0 and 1 − *ϵ* − *κ* ≥ 0. When in a match state, the pair HMM emits letters *b* and *b′* in the *x* and *y* sequences with probability ψ_*b,b′*_; otherwise, in gap states, the probability of letter *b* in the non-gapped sequence is π_*b*_.

Define the MuE transition matrix

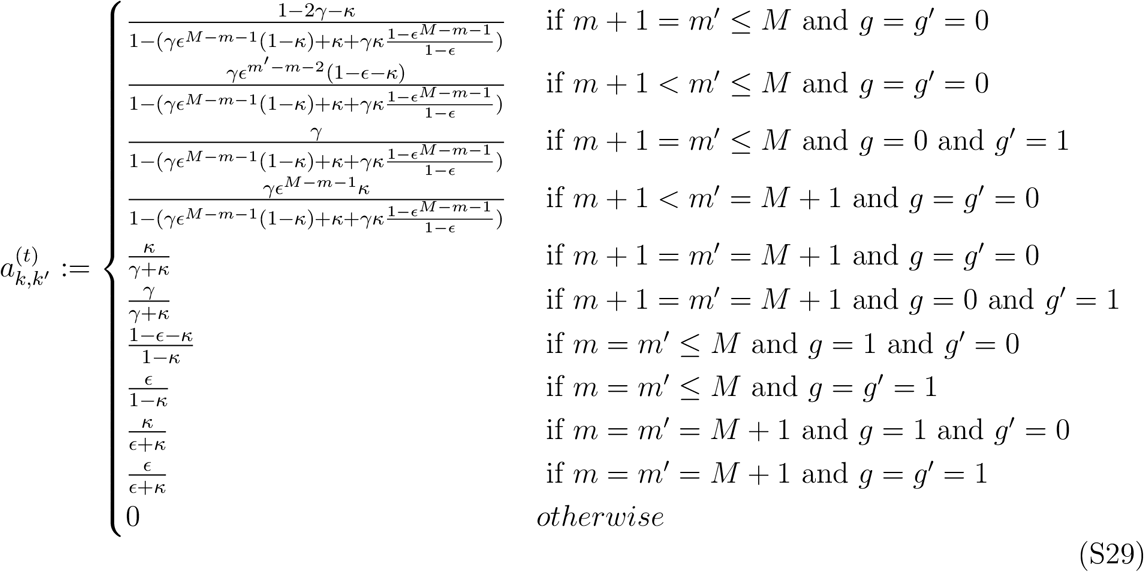

where as before, we refer to states *k* by tuples (*m, g*), where *m* = (*k* + *k*%2)/2 and *g* = *k*%2. The initial transition vector is defined by 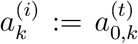. The termination state is *k* = *K* = 2*M* + 2. Define the substitution matrix

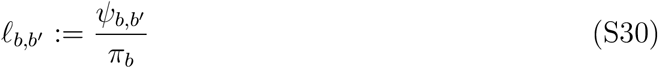

for all *b, b′* ∈ {1, …, *B*}. Let the rows of the insertion matrix *c* be

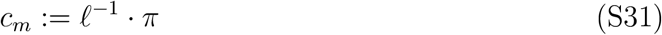

where *ℓ*^−1^ is the inverse of the substitution matrix (assumed to be an invertible matrix). With these definitions, *y* ~ MuE(*x, c, a*, *ℓ*) is equivalent to the conditional distribution of *y* given *x* under the pair HMM.

Note that if γ = 0 then we recover the no-indel case of the MuE distribution (Section S3.2). If, in addition, ψ = *I*_*B*_ then we recover the no-mutation case of the MuE distribution.

**Proof**

We will show that the joint probability of *w* and *y* under the MuE model is identical to the joint probability of the corresponding alignment and *y* under the pair HMM, conditional on *x*. We start by enumerating all possible transitions between states of the MuE Markov chain and computing their probability under the pair HMM model without conditioning on *x*. We use ω^*x*^, ω^*y*^ notation to represent alignments, with the symbol “|” placed to the right of the residue we are transitioning *from*.

1. Transitioning from (*m*, 0) to (*m* + 1 ≤ *M*, 0) has probability 1 − 2γ − *κ*.

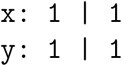
2. Transitioning from (*m*, 0) to (*m′ > m*+1, 0) for *m′ < M* +1 has probability γ*ϵ*^*m′*−*m*−2^(1− *ϵ* − *κ*).

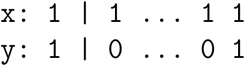
3. Transitioning from (*m*, 0) to (*m* + 1, 1) has probability γ.

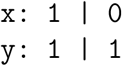
4. Transitioning from (*m < M*, 0) to (*M* + 1, 0), the termination state, has probability γ*ϵ*^*M*−*m*−1^*κ*.

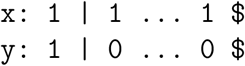
5. Transitioning from (*M*, 0) to (*M* + 1, 0), the termination state, has probability *κ*.

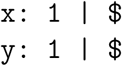
6. Transitioning from (*m*, 1) to (*m* ≤ *M*, 0) has probability 1 − *ϵ* − *κ*.

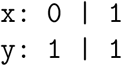
7. Transitioning from (*m*, 1) to (*m*, 1) has probability *ϵ*.

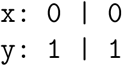
8. Transitioning from (*M* + 1, 1) to (*M* + 1, 0) has probability *κ*

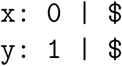
9. Transitioning from the initial state to (1, 0) has probability 1 − 2γ − *κ*.

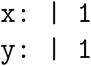
10. Transitioning from the initial state to (*m* > 1, 0) for *m < M* + 1 has probability γ*ϵ*^*m*−2^(1 − *ϵ* − *κ*).

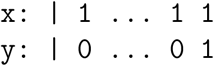
11. Transitioning from the initial state to (1, 1) has probability γ.

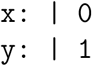
12. Transitioning from the initial state to the termination state has probability γ*ϵ*^*M*−1^*κ* when *M* > 0.

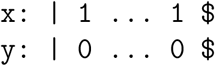
13. Transitioning from the initial state to (*M* +1, 0), the termination state, has probability *κ* when *M* = 0.

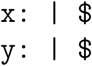

**Figure S13:**
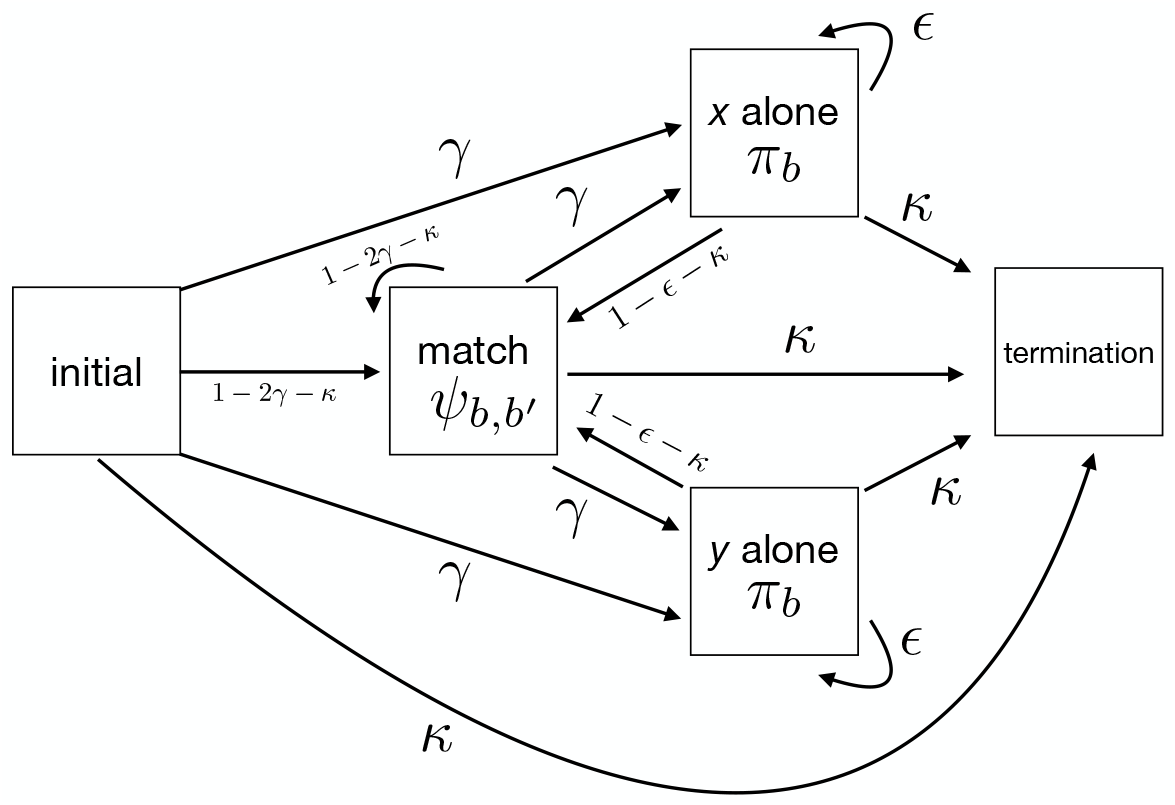
Pair HMM state diagram.

These transition probabilities were derived without conditioning on the fact that *x* has length *M*. To compute this conditional probability, we calculate the probability that the pair HMM generates an alignment with too many or too few *x* residues starting from each MuE Markov model state.

1. Starting from a state (*m < M*, 0), the probability of the pair HMM generating an invalid alignment that is too long (rather than transitioning to a valid MuE state) is γ*ϵ*^*M*−*m*−1^(1 − *ϵ* − *κ*) + γ*ϵ*^*M*−*m*^ = γ*ϵ*^*M*−*m*−1^(1 − *κ*). The first term is from alignments that use a match state instead of terminating.

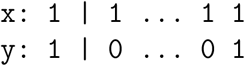 The second term is from alignments that use an *x*-only state instead of terminating.

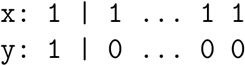
2. Starting from a state (*m < M*, 0), the probability of generating an invalid alignment that is too short (rather than transitioning to a valid MuE state) is 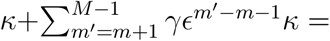 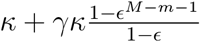. The first term is from alignments that immediately terminate.

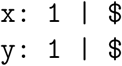 The second term is from alignments that terminate early after transitioning to the *x*-only state.

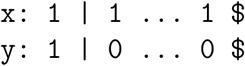
3. Starting from the state (*M*, 0), the probability of generating an invalid alignment is (1 − 2γ − *κ*) + γ = 1 − γ − *κ*. The first term is from alignments that use a match state instead of terminating.

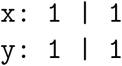 The second term is from alignments that use an *x*-only state instead of terminating.

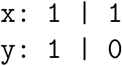
4. Starting from a state (*m ≤ M*, 1) the probability of generating an invalid alignment that is too short is *κ*.

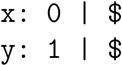
5. Starting from the state (*M* + 1, 1), the probability of generating an invalid alignment that is too long is 1 − *ϵ* − *κ*.

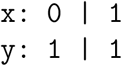
6. Starting from the initial state, the probability of generating an invalid alignment that is too long is γ*ϵ*^*M*−1^(1 − *ϵ* − *κ*) + γ*ϵ*^*M*−*m*^ = γ*ϵ*^*M*−1^(1 − *κ*). The first term is from alignments that use a match state instead of terminating.

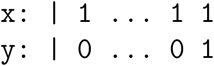 The second term is from alignments that use an *x*-only state instead of terminating.

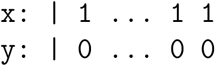
7. Starting from the initial state, the probability of generating an invalid alignment that is too short is 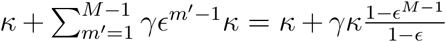 when *M* > 0. The first term is from alignments that immediately terminate.

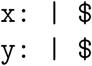 The second term is from alignments that terminate early after transitioning to the *x*-only state.

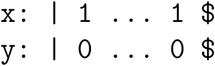
8. Starting from the initial state, if *M* = 0, then the probability of generating an invalid alignment is (1 − 2γ − *κ*) + γ = 1 − γ − *κ*. The first term is from alignments that use a match state.

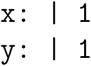 The second term is from alignments that use an *x*-only state.

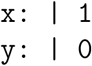

We can confirm that all possible trajectories of the pair HMM are either valid transitions under the MuE Markov model or produce alignments with too few or too many *x* residues, by checking that the outward transition probabilities from each state sum to one.

1. From a state (*m < M*, 0), the total outward transition probability is

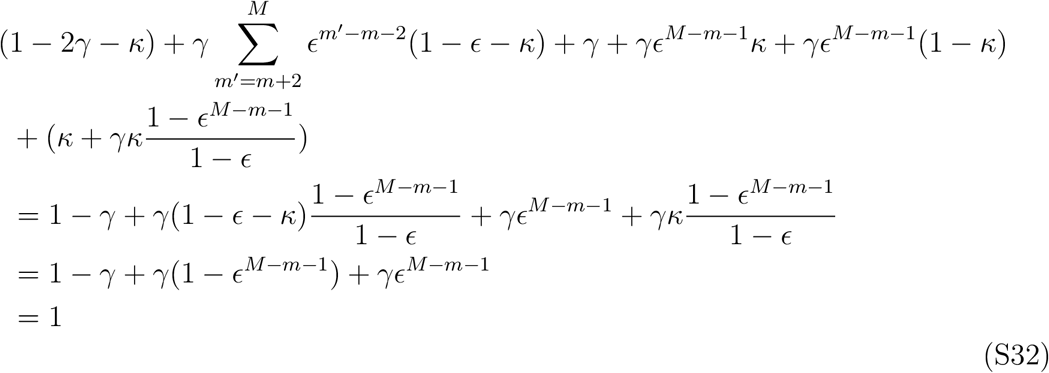
2. From the state (*M*, 0), the total outward transition probability is

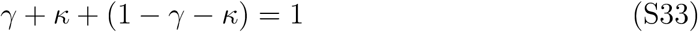
3. From a state (*m* ≤ *M*, 1), the total outward transition probability is

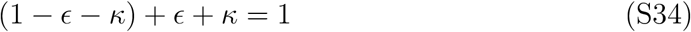
4. From the state (*M* + 1, 1), the total outward transition probability is

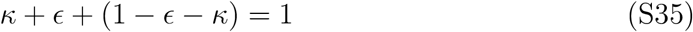
5. From the initial state, with *M* > 0, the total outward transition probability is

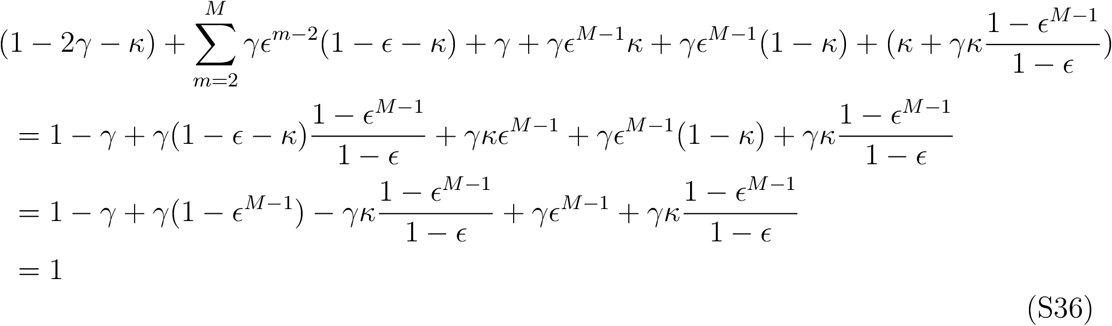
6. From the initial state, with *M* = 0, the total outward transition probability is

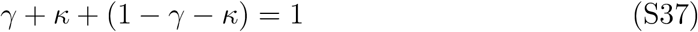

Consolidating transition probabilities and conditioning on the length of *x* yields the transition matrix Equation S29.

Next we consider sequence emission probabilities, given an alignment. Recall that *x* and *y* are one-hot encodings of sequences.

1. Consider the case that *y*_*l*_ is aligned to *x*_*m*_, ie.

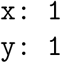 The conditional probability of *y*_*l,b*′_ = 1 given *x_m,b_* = 1 is, according to the pair HMM, ψ_*b,b′*_ /π_*b*_. This matches the conditional probability assigned by the MuE,

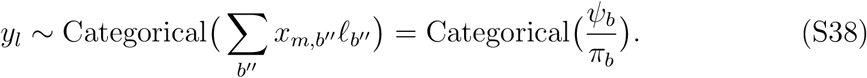
2. Consider the case that *y*_*l*_ is aligned to a gap, ie.

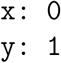 The conditional probability of *y_l,b_* given *x* is just π_*b*_ (since *x* is not informative in this case). This matches the conditional probability assigned by the MuE,

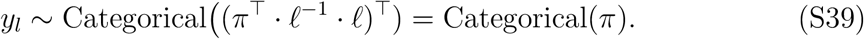

where ⊤ is the transpose symbol.
3. Consider the case that *x*_*m*_ is aligned to a gap, ie.

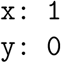 The conditional probability of *x*_*m*_ given *x* is trivially one, so this term does not contribute to the conditional probability of *y* given *x* under the pair HMM. It also does not contribute to the probability under the MuE.

Thus, term-by-term, the joint probability of *w* and *y* under the proposed MuE distribution matches the joint probability of the corresponding alignment and *y* under the pair HMM conditional on *x*.

#### S4.4 Profile HMM

The profile HMM (pHMM) is a widely used model for defining protein sequence families, inferring multiple sequence alignments, and performing database searches [16]. Define the pHMM insertion parameter *r_m,j_* ∈ [0, 1] for all *m* ∈ {1, …, *M* + 1} and *j* ∈ {0, 1, 2}, and the deletion parameter *u_m,j_* ∈ [0, 1] for all *m* ∈ {1, …, *M* + 1} and *j* ∈ {0, 1, 2}, with *u*_*M*+1*,j*_ = 0 for *j* ∈ {0, 1, 2}. Then define the MuE transition matrix

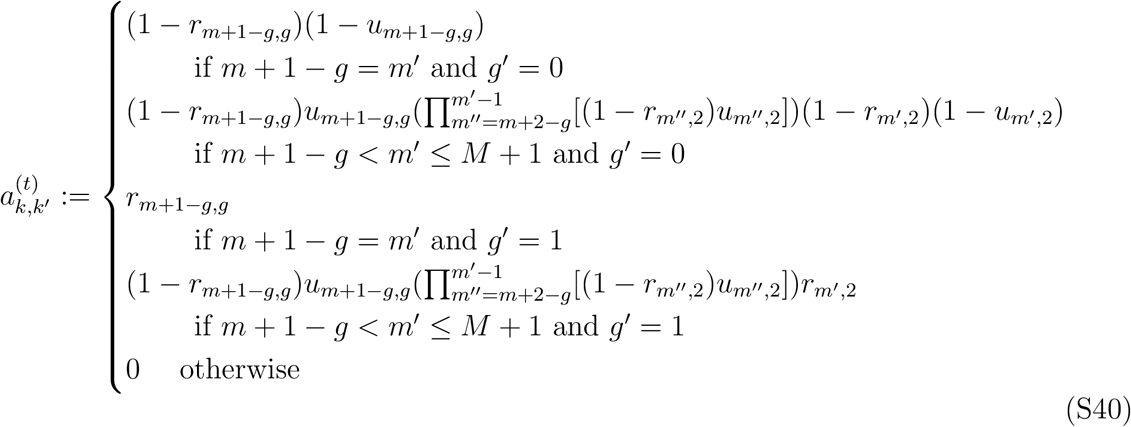

where as before, we refer to states *k* by tuples (*m, g*), with *m* = (*k* + *k*%2)/2 and *g* = *k*%2. The initial transition vector is defined by 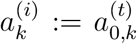. The state *k* = *K* = 2*M* + 2 is the termination state. Let the MuE substitution matrix *ℓ* be the identity matrix *I*_*B*_, ie.

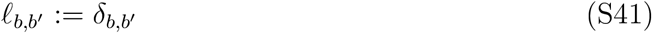

for *b, b′* ∈ {1, …, *B*}. Then the profile HMM can be written as

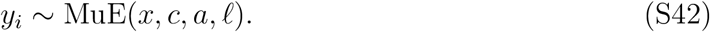

**Figure S14:**
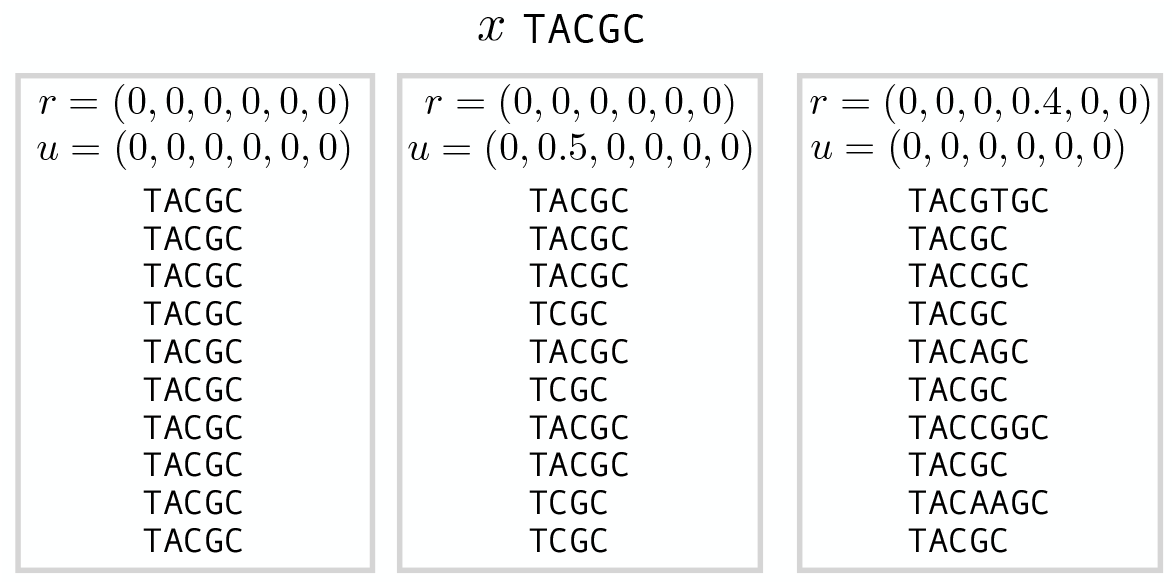
Samples from the profile HMM. The “ancestral” sequence *x* is set to `TACGC`, and we set *r_m,j_*_=0_ = *r_m,j_*_=1_ = *r_m,j_*_=2_ and *u_m,j_*_=0_ = *u_m,j_*_=1_ = *u_m,j_*_=2_ for all *m*.

Figure S14 illustrates samples from the pHMM. Intuitively, *r* controls insertion probabilities and *u* controls deletion probabilities; when *r_m,j_* = 0 and *u_m,j_* = 0 for all *m* and *j*, we recover the no-mutation case of the MuE, since *ℓ* is the identity matrix (Section S3.2).

**Proof**

This result follows from the relabeling of the profile HMM Markov state architecture with the (*m, g*) notation used for the MuE distribution (Figure S15). “Deletion states” in a profile HMM do not generate observations *y*_*l*_. To compute the probability of transitioning between two observable states (*m, g*) and (*m′*, *g′*), we compute the probability of (1) direct paths between the two states and (2) all possible paths between the two states that go only through deletion states. This yields Equation S40.

**Figure S15:**
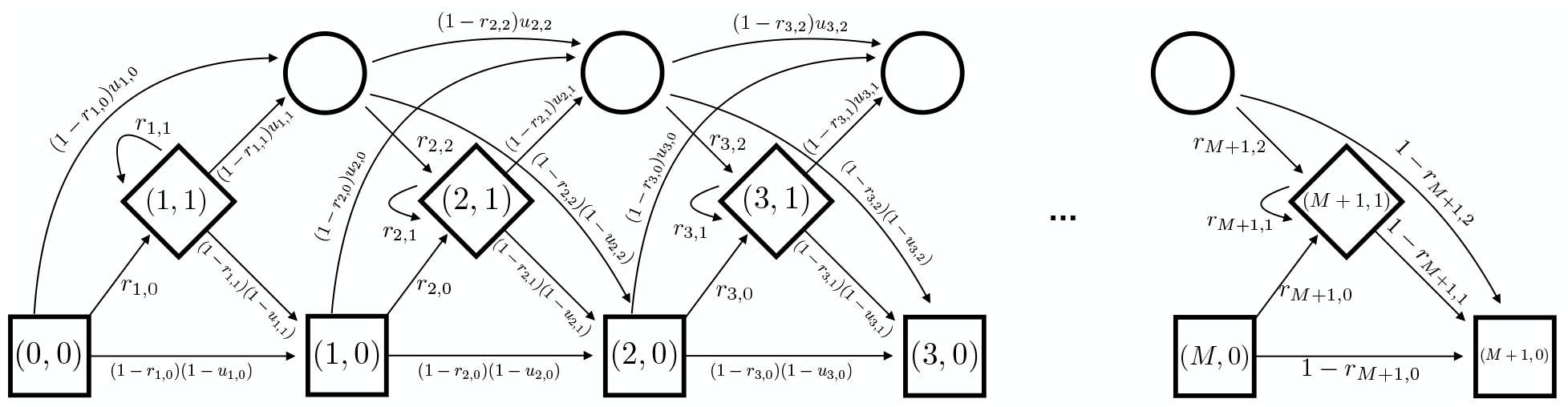
Profile HMM state architecture. The conventional profile HMM state architecture labeled with MuE states, using (*m, g*) notation; recall that *k* = 2*m − g*. The state (0, 0) represents the initial state, and the state (*M* + 1, 0) represents the termination state. Squares indicate match states, diamonds indicate insertion states, and circles indicate “deletion states”.

The emission probability of each state in the pHMM is set by its associated emission vector. Without loss of generality, we can write any emission matrix of the pHMM as *e* = ξ · *x* + ζ · *c*. This is equivalent to the MuE emission matrix, since *ℓ* is set to the identity matrix.

#### S4.5 Needleman-Wunsch

The Needleman-Wunsch (NW) algorithm is a classic non-probabilistic alignment method [48]. Let *G* be the NW gap penalty (assumed to be negative) and define *u* := *e*^*G*^. We define the MuE transition matrix

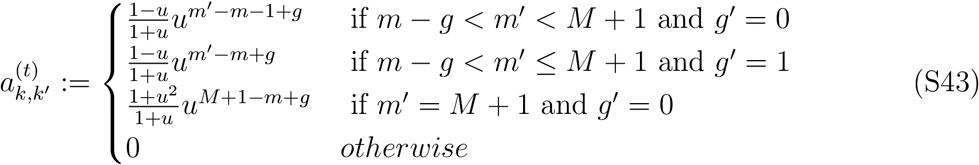

where as before, we refer to states *k* by tuples (*m, g*), where *m* = (*k* + *k*%2)/2 and *g* = *k*%2. The initial transition vector is defined by 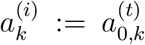. The state *k* = *K* = 2*M* + 2 is the termination state. Let *S*_*b,b*′_ be the NW similarity matrix, for which we assume that 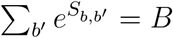 for all *b*. We define, for *b, b′* ∈ {1, …, *B*},

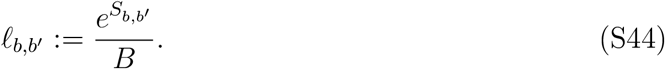

Finally, for all *m* ∈ {1, …, *M* + 1},

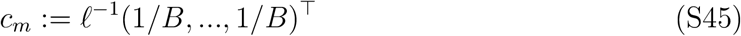

where *ℓ*^−1^ is the inverse of the substitution matrix (assumed to be invertible) and (1*/B, …*, 1*/B*)^⊤^ is a length *B* vector. Let *x* and *y* each be one-hot sequence encodings. Now, under the MuE model *y* ~ MuE(*x, c, a*, *ℓ*), the maximum *a posteriori* estimator of the alignment variable *w* given *x* and *y* corresponds to the Needleman-Wunsch pairwise alignment between *x* and *y*.

Note that in the limit *G* → −∞ we recover the no-indel case of the MuE distribution. If, in addition, *S*_*b,b*′_ → −∞ for all *b′* ≠ *b*, we recover the no-mutation case of the MuE distribution.

**Proof**

We can organize the NW scoring system according to transitions in the MuE Markov model. We use ω^*x*^, ω^*y*^ notation to represent alignments, with the symbol “|” placed to the right of the residue we are transitioning *from*. We assign *l′* to be the residue of *y* at the column of the alignment corresponding to state *k′*.

1. Transitioning from (*m*, 0) to (*m′ > m*, 0) gives a NW score of (*m′* − *m* − 1)*G* + ∑*_b,b′_x_m′,b_S_b,b′_y_l′,b′_*.

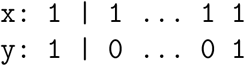
2. Transitioning from (*m*, 0) to (*m′ > m*, 1) gives a NW score of (*m′* − *m*)*G*

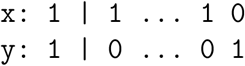
3. Transitioning from (*m*, 1) to (*m′* ≥ *m*, 0) gives a NW score of (*m′*−*m*)*G*+∑_*b,b′*_*x*_*m′,b*_*S_b,b′_y*_*l′,b′*_

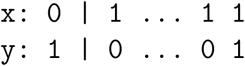
4. Transitioning from (*m*, 1) to (*m′* ≥ *m*, 1) gives a NW score of (*m′* − *m* + 1)*G*.

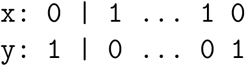
5. Transitioning from (*m*, 0) to (*M* + 1, 0) gives a NW score of (*M* − *m* − 1)*G*.

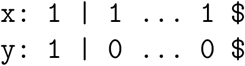
6. Transitioning from (*m*, 1) to (*M* + 1, 0) gives a NW score of (*M* − *m*)*G*.

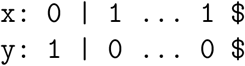

Now we can rewrite the Needleman-Wunsch objective function in terms of these transitions, rather than in terms of gap and insert scoring. In particular, define

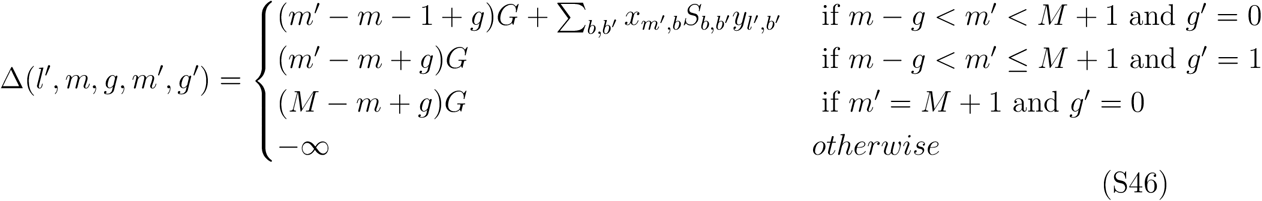

Based on the cases outlined above, the NW objective function can now be rewritten as

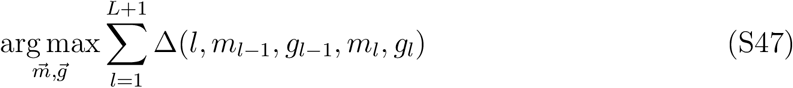

where the vectors 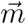 and 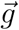 are each length *L* + 2 and we set *m*_0_ = 0, *g*_0_ = 0, *m*_*L*+1_ = *M* + 1, *g*_*L*+1_ = 0. If we find the solution to this objective function, then follow the mapping from the list of Markov chain states (*m*_1_, *g*_1_), *…*, (*m*_*L*+1_, *g*_*L*+1_) back to an alignment (Section S4.1.3), we obtain the Needleman-Wunsch alignment between sequences *x* and *y*.

Now we examine the maximum *a posteriori* estimator of *w* under the MuE distribution.

We have

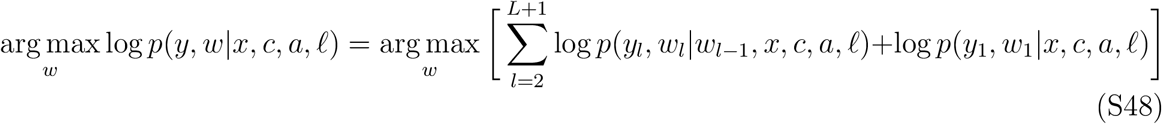

Let *k*_*l*_ = arg max*_k_ w_l_* be the state of the Markov model at the *l*th residue and let *m*_*l*_ = (*k*_*l*_ + *k*_*l*_%2)/2 and *g*_*l*_ = *k*_*l*_%2. We then have, under the MuE model,

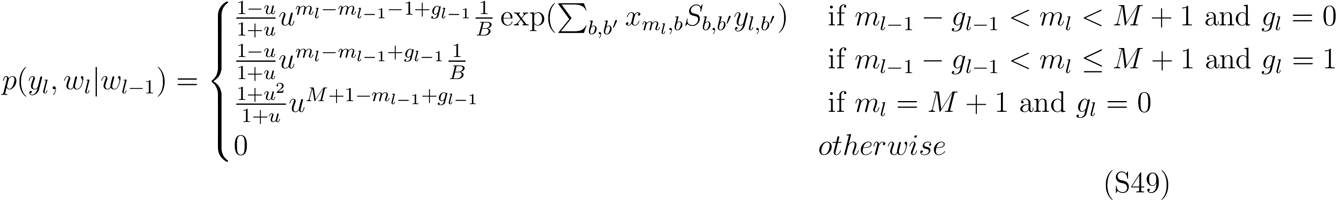

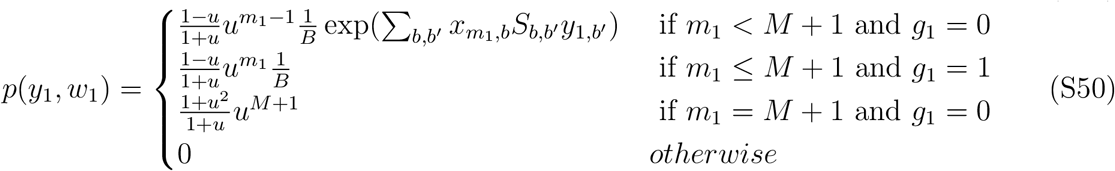

Note that we must have *m*_*L*+1_ = *M* + 1 and *g*_*L*+1_ = 0, since the sequence *y* is of length *L*. Now, the maximum *a posteriori* estimator of *w* can be written as

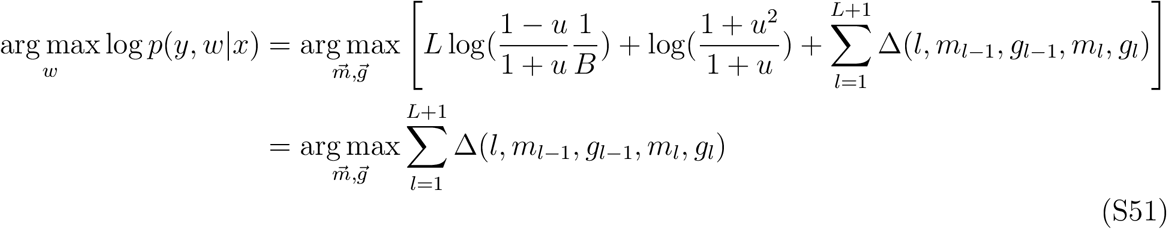

where *m*_0_ = 0 and *g*_0_ = 0. This objective function is identical to the NW objective function (Equation S47), so the maximum *a posteriori* estimator of *w* in the MuE distribution corresponds to the Needleman-Wunsch pairwise alignment of *x* and *y*.

#### S4.6 Vogel et al. Natural Language Translation

An interesting point of comparison for the MuE distribution is the natural language translation model presented in Vogel et al. [49]. This model takes the same form as a MuE distribution, with *x* a one-hot encoding of a sentence in one language and *y* one-hot encoding of a sentence in another language. With states *k* indexed by tuples (*m, g*), where *m* = (*k* + *k*%2)/2 and *g* = *k*%2, the transition matrix takes the form

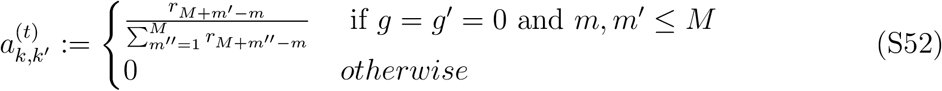

where 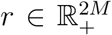 is a vector of non-zero weights. The initial transition vector is defined by 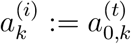. The length of *y* is sampled independently of *w*.

However, the Vogel et al. model is not a MuE distribution: the transition matrix does not satisfy the condition that 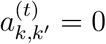 whenever *k′* + *k′*%2 − *k* + *k*%2 ≤ 0 for all accessible states *k*, so the Vogel et al. model does not produce valid biological sequence alignments.

### S5 Model Details

In this section we provide a detailed description of the models evaluated in the main text. For the MuE distribution in each, we choose *a* to fit the form of the profile HMM (Section S4.4) with the additional restrictions *r_m,j_*_=0_ = *r_m,j_*_=1_ = *r_m,j_*_=2_ =: *r*_*m*_ and *u_m,j_*_=0_ = *u_m,j_*_=1_ = *u_m,j_*_=2_ =: *u*_*m*_. We set *u*_*M*+1_ = 0. Intuitively, *r*_*m*_ is the probability of an insertion at position *m* of *x* and *u*_*m*_ is the probability of a deletion at position *m* of *x*. The transition matrix is

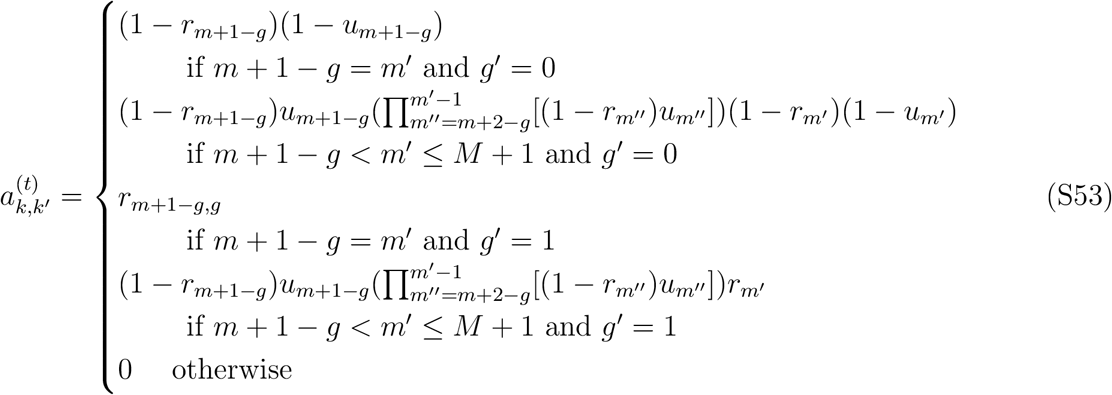

where, as in Section S4.4, *m* = (*k*+*k*%2)/2, *g* = *k*%2, *m′* = (*k′*+*k′*%2)/2 and *g′* = *k′*%2. The initial transition vector follows the same form as the transition matrix, and can be written as 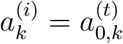. Rather than assign a termination state we assume the length of the sequence *y*_*i*_, ie. *L*_*i*_, is independent of *w* (Section S3.1); for convenience, we assign *p*(*y*_*l*_|*w_l,K_* = 1) = 0 for all possible *y*_*l*_, making the state *k* = *K* inaccessible. Since the probability of *L*_*i*_ does not contribute to the per residue perplexity performance metric (Section S7) we do not use an explicit model for *L*_*i*_.

Note that in our experiments we go slightly beyond the vanilla H-MuE presented in the main text (Equation 2), and allow the insertion sequence *c* to depend on the continuous-space vector model *p*(*v*|*θ*).

#### S5.1 Profile HMM

The profile HMM is

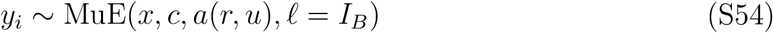

where *a*(*r, u*) depends deterministically on the parameters *r* and *u* according to Equation S53, *D* = *B* and *I*_*B*_ is the *B* × *B* identity matrix.

#### S5.2 RegressMuE

The RegressMuE model uses a linear regression model as the H-MuE’s continuous-space vector model. Let *h_i,_*_1_, …, *h_i,T_* be covariates associated with sequence *y*_*i*_. Let 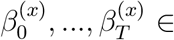 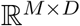 be a set of coefficients associated with *x*, and let 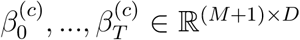 be a set of coefficients associated with *c*. Then the RegressMuE is

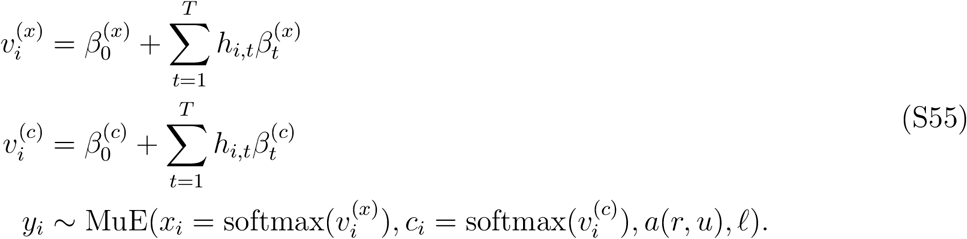

Note that in this model, unlike the pHMM, the substitution matrix *ℓ* is not constrained to the identity. In the no-indel special case of the MuE (Section S3.2), when *r*_*m*_ = *q*_*m*_ = 0 for all *m* and *ℓ* = *I*_*B*_, the RegressMuE reduces to a multi-output multinomial logit regression model.

#### S5.3 FactorMuE

The FactorMuE model is the latent linear version of the RegressMuE. Instead of observing covariates *h*, we draw a latent variable *z* from a standard normal prior,

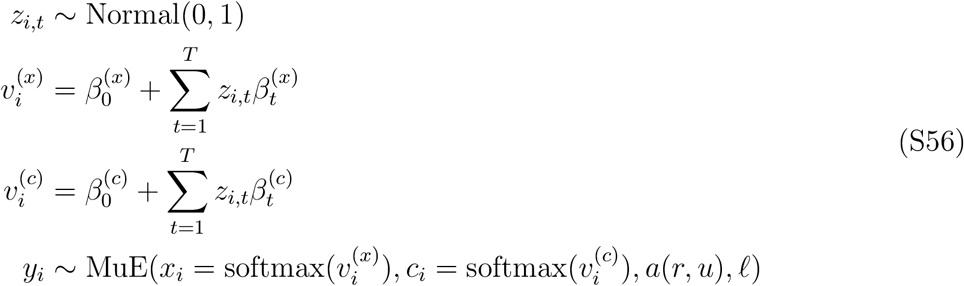

#### S5.4 NeuralMuE

The NeuralMuE model uses a fully connected neural network as the H-MuE’s continuous-space vector model. We use a Γ-layer network with relu nonlinearities, widths *T*_1:(Γ+1)_, and weights *β*_1:(Γ+1)_. Let 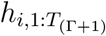 be a vector of covariates.

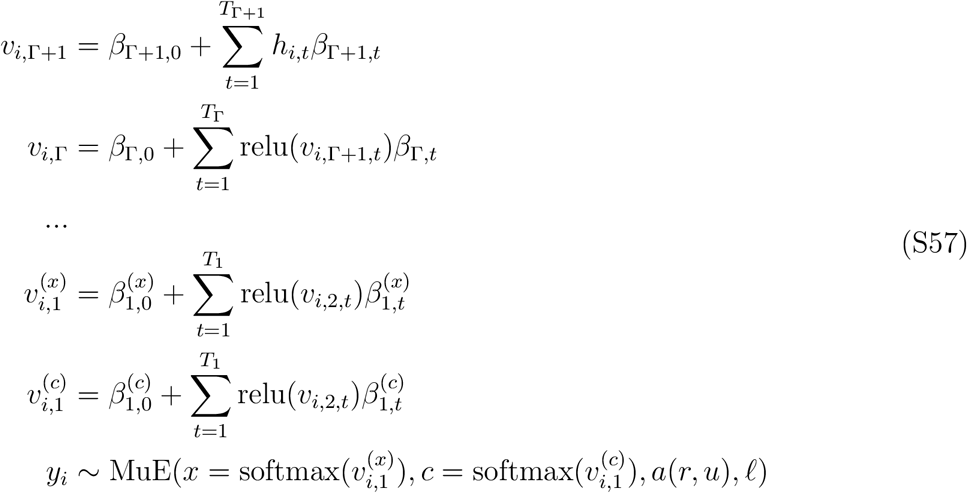

#### S5.5 LatentNeuralMuE

The LatentNeuralMuE model uses a neural network latent variable model as the H-MuE’s continuous-space vector model. It is the latent covariate version of the NeuralMuE, where instead of observing *h* we draw a latent variable *z* from a standard normal prior.

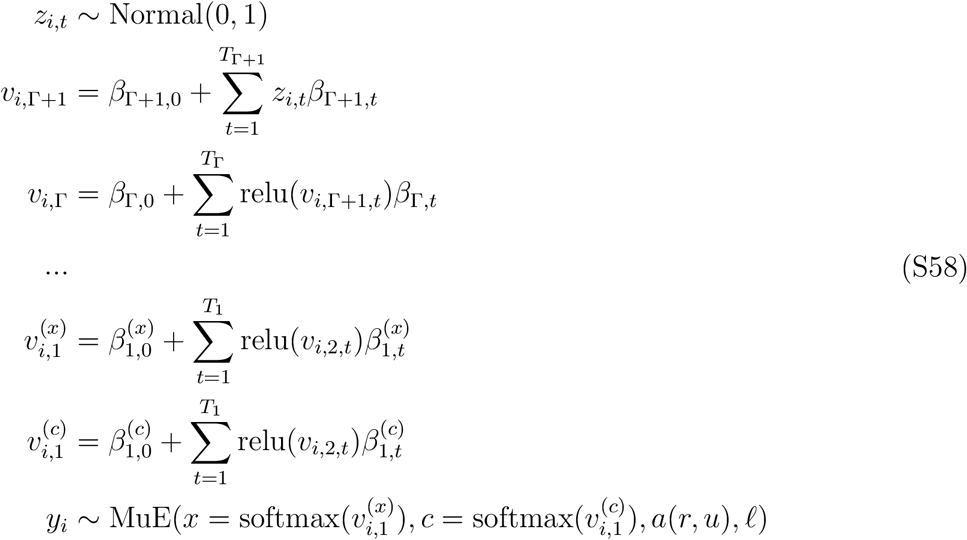

#### S5.6 Priors

We place standard normal priors Normal(0, 1) over each element of each coefficient matrix *β* in each model. Recall that each row of the matrix *ℓ* is constrained to the simplex, *ℓ* ∈ Δ_*B*−1_. To enable easy gradient-based optimization and stochastic variational inference [8], we transform an unconstrained parameter 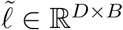 with a Gaussian prior to the simplex,

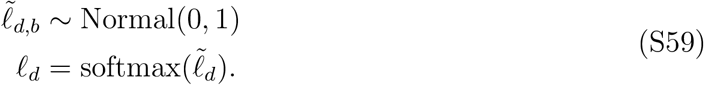

The variables *r*_*m*_ and *u*_*m*_ are constrained to [0, 1]. This corresponds to the first dimension of a simplex Δ_1_, and so we apply the same approach,

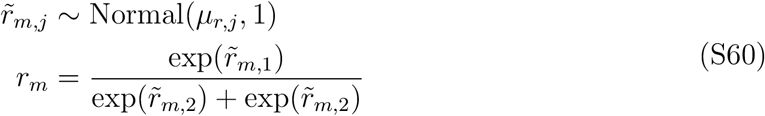

for *j* ∈ {1, 2}. The variable *u*_*m*_ is handled identically, with prior 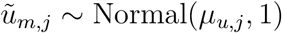.

### S6 Inference

Variational inference approximates the posterior distribution *p*(*θ*|*y*_1:*N*_) of a given probabilistic model using a tractable family of distributions *q*_η_(*θ y*_1:*N*_) parameterized by η [10]. To form this approximation, variational inference minimizes the Kullback-Leibler (kl) divergence between the two distributions,

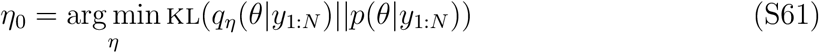

This objective can be rewritten as maximizing the evidence lower bound (elbo),

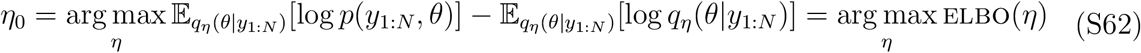

We employ mean-field variational inference for H-MuE models. We use a diagonal Gaussian distribution, with unknown mean and standard deviation, for the variational distribution over the global parameters 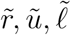 and 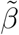. For the local variable *z* in the FactorMuE and LatentNeuralMuE, we amortize inference using a recognition network (an encoder) [12, 13]. In particular, we set

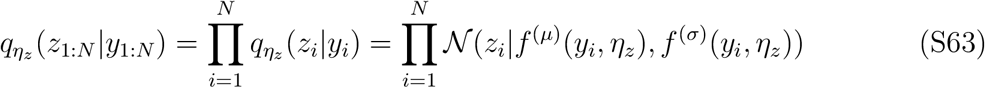

where 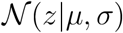 is the probability distribution function of a Gaussian with mean *μ* and standard deviation σ, and *f* ^(*μ*)^(*y_i_*, η_*z*_) and *f* ^(σ)^(*y_i_*, η_*z*_) are differentiable functions of η_*z*_. We parameterize *f* ^(*μ*)^ and *f* ^(σ)^ using a neural network,

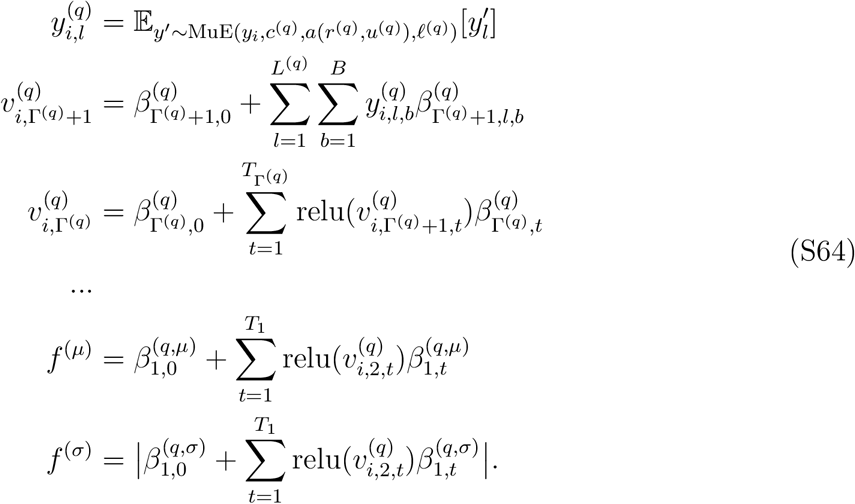

where we have introduced the variational parameters (*β*^(*q*)^, *c*^(*q*)^, *r*^(*q*)^, *u*^(*q*)^, *ℓ*^(*q*)^) = η_*z*_. The first layer of the encoder employs the MuE distribution and computes the expected value of mutants of *y*_*i*_, at positions *l* ∈ {1, …, *L*^(*q*)^}; this expected value is a differentiable function of the MuE parameters, and can be tractably computed using the forward algorithm. We use the same parameterization of the MuE distribution as in the model (Section S5), but fix 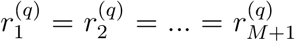 and 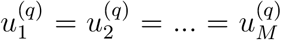 and 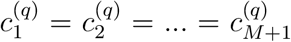. Intuitively, the MuE encoding serves to “smear out” the one-hot encoded sequence *y*_*i*_ according to learnable indel and substitution probabilities, making it easier for the encoder to learn which sequences are similar, and making each encoded sequence 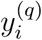 the same length *L*^(*q*)^.

To optimize the variational approximation we need to compute the gradient of the ELBO with respect to the variational parameters η. To enable faster optimization we employ stochastic variational inference, approximating the gradient at each update step using a minibatch of data [50]. Let *ϕ* = (*β*, *r, u*, *ℓ*) be the global parameters of the H-MuE models and let η_*ϕ*_ be the parameters of the associated mean-field variational distribution. Then the gradient of the ELBO is

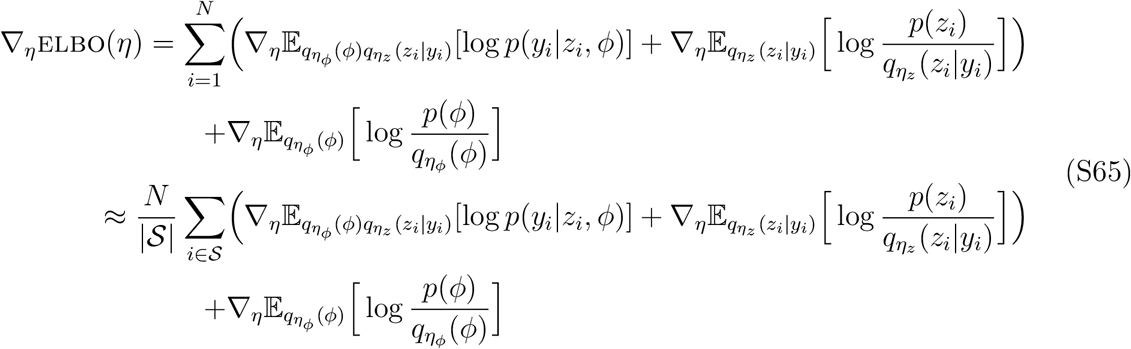

where 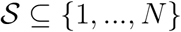 is the set of datapoint indices making up the minibatch and 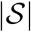 is the size of the set 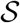. We estimate the gradient of the first term on the right hand side of this equation using the reparameterization trick Monte Carlo estimator (with a single sample) and automatic differentiation [8, 12, 13]. The remaining terms can be computed analytically (see e.g. [12, 13]). Note that this approach relies crucially on the fact that the marginal likelihood of the MuE model, *p*_MuE_(*y|x, c, a*, *ℓ*) = ∑_*w*_ *p*_MuE_(*y|w, x, c, a*, *ℓ*), is a differentiable function of *x*, *c*, *a* and *ℓ*. We integrate over all possible values of the Markov chain state variable *w* using the forward algorithm.

It is useful in some circumstances to reweight the variational objective to reduce the amount of regularization placed on the local latent variable. In particular, for χ ∈ [0, 1], we reweight the ELBO as

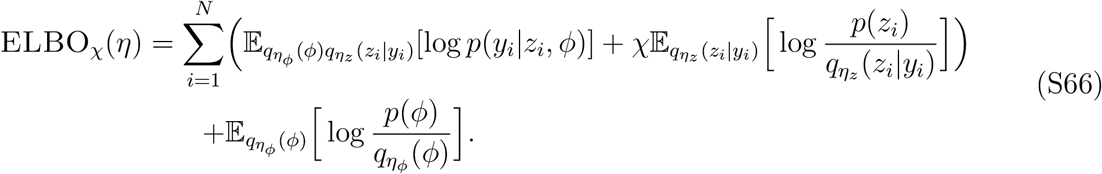

We achieved improved training performance by annealing the weight χ from 0 to 1 linearly over the course of an initial time period during training [51]. To avoid posterior collapse and produce informative latent representations, we found it useful in certain cases to anneal χ only up to a low value χ_0_ < 1; this annealing schedule was only used for producing data visualizations, rather than prediction of held out data (Section S10) [52].

### S7 Perplexity

The per residue perplexity of a probabilistic sequence model *p*(*y*), over a dataset *y*_1:*N*_, is defined as

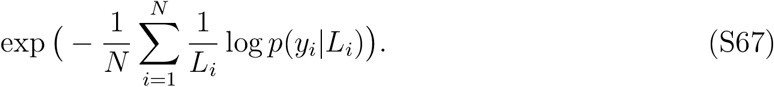

In evaluating our models, we computed the average log likelihood performance on a heldout test set 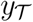 for the ensemble of models learned from the training set 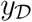. More precisely, we use

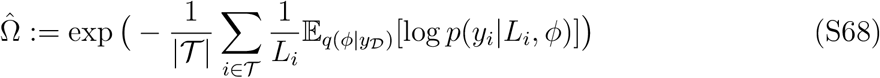

where 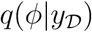 is the variational approximation to the posterior distribution from the training dataset and 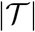 is the size of the test set. For models with local latent variables *z*_*i*_, we approximate the marginal likelihood using the ELBO [10],

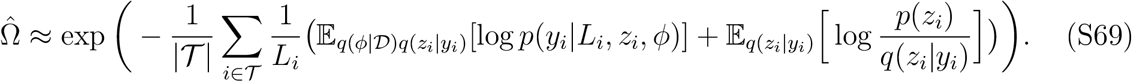

We use Monte Carlo estimation for the expectations. In comparing between different models *p*_1_ and *p*_2_, we also report the log Bayes factor associated with the held out data, ie. the difference in total log probability of the heldout data between the two models,

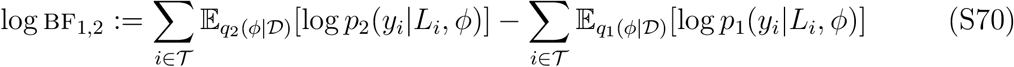

where *q*_1_ and *q*_2_ are the variational approximations associated with *p*_1_ and *p*_2_. For models with local latent variables, we can use the ELBO approximation as in Equation S69. The Bayes factor provides a measurement of the total evidence in favor of one model versus another.

Per residue perplexity is a useful performance metric for biological sequence models because it is an absolute scale and comparable across datasets as well as models. In the interest of making this scale interpretable, we computed the expected per-residue perplexity for a variety of different protein sequence models, covering different regimes. In particular, for each model *p*(*y*), we examined the expected perplexity in the large data limit, assuming that the model is true,

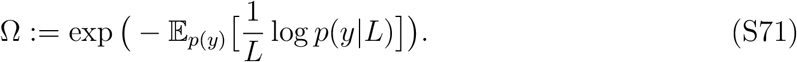

The expected perplexity is the exponentiated entropy of the model distribution, and so provides a measurement of sequence diversity under the model. Below, we compute the expected perplexity for distributions ranging from the very high diversity regime (all of evolution) down to the very small diversity regime (human population genetics).

#### Naive

A naive model assigns an equal probability to each amino acid. In this case the per residue perplexity is

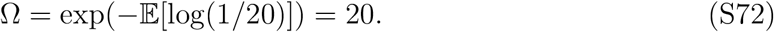

#### Amino acid frequencies

A simple modeling approach is to predict individual amino acids solely based on their naturally occurring frequency across evolution. Using the UniprotKB amino acid frequencies *f*_*b*_ for *b* ∈ {1, …, *B* = 20}, we have

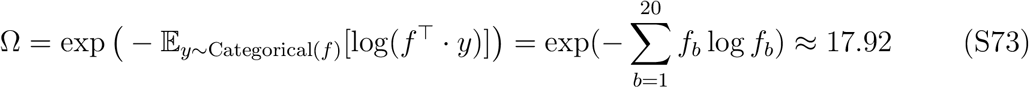

where *y* is a one-hot encoding of an amino acid and ⊤ is the vector transpose symbol [53, 54].

#### BLOSUM62

If we are studying specific evolutionary families of proteins, an idealized strategy for building a model is to infer the sequence of the last common ancestor and then predict family members using the standard BLOSUM62 substitution matrix [55]. The BLOSUM62 matrix is a renormalized copula density, but we can convert it into a mutation probability matrix *ℓ* by assuming the marginal probability of each amino acid follows the UniprotKB frequency across evolution:

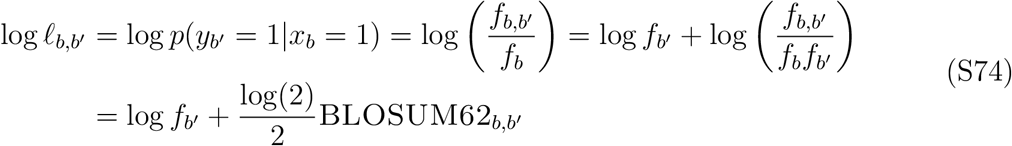

where *x* is a one-hot encoding of the ancestral amino acid, *y* is a one-hot encoding the mutated amino acid, and *f*_*b,b*′_ is the joint probability of amino acids *b* and *b′*, where *b, b′* ∈ {1, …, *B* = 20}. (The log(2)/2 factor comes from the definition of BLOSUM62.) We renormalize the rows *ℓ*_*b*_ to ensure *ℓ*_*b*_ ∈ Δ_*B*−1_ (BLOSUM62 uses only small integers, producing non-negligible rounding error). Next, we assume that the ancestral sequence is known exactly, has infinite length, and the distribution of each amino acid within the ancestral sequence follows the UniprotKB overall frequency across evolution. The expected per residue perplexity is

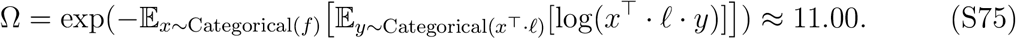

#### Human Population Genetics

Finally, we examined a simple model of human population variation. Each human has on average roughly 5 million single nucleotide polymorphisms (SNPs) relative to the reference genome [56]. Naively assuming a constant mutation rate over the genome, the probability of a mutation occurring in any particular codon is *q*_codon_ = 1 − (1 − 5/6400)^3^, since there are 6.4 billion total base pairs. If we very naively assume a uniform probability of the codon mutating to any other amino acid, then we can use the substitution matrix *ℓ* defined by

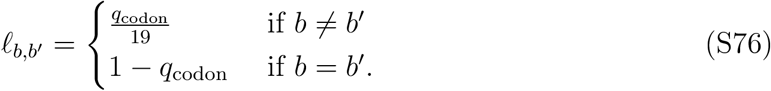

If we further naively assume that there are no correlations among mutations at different genome locations when looking across individuals, then the expected per residue perplexity of the sequence distribution is

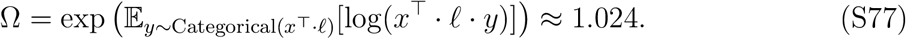

### S8 Density Estimation

Evolutionarily related sequences were collected using jackhmmer (v3.1) from the UniRef100 dataset (date 6/2019) [57, 58]. We used seed sequences with Uniprot identifiers DYR HUMAN (DHFR dataset), PINE ECOLI (PINE dataset), CDN1B HUMAN (CDKN1B dataset), and VE6 HPV16 (VE6 dataset). We set a bitscore threshold of 0.5 bits/residue as in [17] and ran the jackhmmer search using the API from the EVcouplings package [59]. We included the full envelope of the profile HMM hit (ie. including residues classified as insertions and deleting gap symbols) in the final dataset. The CDN1B dataset had 1,055 sequences and the VE6 dataset 1,609 sequences. We found 32,510 and 79,354 hits respectively for the DHFR and PINE datasets, which we randomly subsampled to 10,000 sequences to create the final datasets. Note that the jackhmmer search algorithm uses a profile HMM to find distant homologs, and thus may bias the dataset to look more like samples from a pHMM; we therefore expect the performance gains from using H-MuE models, as compared to the pHMM, on these datasets to be smaller (more conservative) than the performance gains that might be achieved on alternative datasets assembled using different search methods. The TCR dataset was not assembled using jackhmmer (Section S9).

We set the latent alphabet size *D* = 25. In each experiment, we set *M* to be 10% longer than the longest sequence in the dataset. We used *T* = 5 latent space dimensions in the FactorMuE and layer sizes *T*_2_ = 5, *T*_1_ = 10 in the LatentNeuralMuE (we found a substantial dropoff in performance when increasing network width or depth). In the recognition network, we set *L*^(*q*)^ = *M*. We also used Γ^(*q*)^ = 0 (no relu nonlinearities) in the FactorMuE recognition network and Γ^(*q*)^ = 1, *T*_1_ = 10 in the LatentNeuralMuE recognition network. For the prior on the MuE insertion and deletion parameters we used *μ_r_* = *μ_u_* = (100, 1) to disfavor indels.

We optimized the variational approximation using Adam [60] and a batch size of 5. The mean of the variational distribution was initialized at the prior mean, while the variance was initialized to a small random value (the absolute value of a sample from a normal distribution with standard deviation 0.01). We used one Monte Carlo sample to estimate the ELBO gradient at each step. For each model and dataset, we evaluated two different learning rates, 0.1 and 0.01, and three different random restarts, selecting among training runs the parameter values that reached the highest ELBO on the training set for making predictions. For models with local latent variables (the FactorMuE and LatentNeuralMuE), we annealed the ELBO reweighting factor χ from 0 to 1 linearly over the first 2 epochs. We trained for 4 epochs total on the DHFR and PINE datasets, and 7 epochs total on the smaller CDKN1B, VE6 and TCR datasets, which was sufficient for convergence in each model. We estimated the heldout perplexity using one independent Monte Carlo sample per batch. Computations were performed on graphics processing units (GPUs), and we used gradient accumulation to reduce memory usage.

### S9 T-Cell Receptor Analysis

We downloaded a publicly available dataset of 6,327 T-cell receptor (TCR) sequences found in CD8+ cytotoxic T-cells https://support.10xgenomics.com/single-cell-vdj/datasets/2.2.0/vdj_v1_hs_cd8_t [35]. These were sequenced using single cell sequencing of peripheral blood mononuclear cells obtained from an individual healthy donor. Internal stop codons were deleted from the sequence. We used the provided CellRanger annotations of chain features.

To obtain a latent space representation (Figure 3BCD), we trained the FactorH-MuE model with *T* = 2 latent dimensions, and chose among training runs based on a randomly held out test set (5% of the data). Hyperparameters were otherwise set as in Section S8.

We computed the magnitude of the shift in sequence space along a vector with tail *z*_0_ and head *z*_1_ as

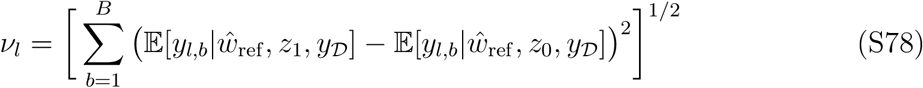

where 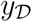 is the training dataset. The expectation is estimated using the variational approximation to the posterior of the FactorMuE (using 10 Monte Carlo samples). 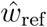 is the maximum *a posteriori* estimator of *w* given the sequence *y*_ref_ of the reference protein structure PDB:2BNR (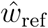 is estimated using a single sample from the variational approximation to the posterior and the Viterbi algorithm). Fixing a particular *w* value enables easy visual comparison between “aligned” sequences position-by-position.

To understand how the full TCR sequence differs between *α* and *β* chains we used the RegressMuE, with the covariate vector *h*_*i*_ a one-hot encoding of the chain type annotated by CellRanger; sequences without an annotation were encoded as (0, 0). We computed the regression shift *ν*_*l*_ in the same way as Equation S78, with the covariate *h* in place of *z*.

### S10 Influenza Analysis

We downloaded publicly available influenza A(H3N2) HA sequences from GISAID [26]. We selected only sequences longer than 500 amino acids and with no ambiguous amino acids. Some sequences were labeled at different levels of time resolution, with annotations providing months or years rather than days; we assumed month and/or day were missing at random and imputed them uniformly at random. Following Lee et al. [42], we randomly subsampled six sequences per month, from 1968 to October 2019, to form the dataset. In the forecasting experiments we removed the mis-annotated data identified in the 2008 outlier cluster marked by ‡ in Figure 5 prior to subsampling (GISAID identifiers EPI_ISL_24813, EPI_ISL_24814, …, EPI_ISL_24867). Our main results were stable upon resampling. We extracted only the first 350 amino acids of each HA sequence, covering HA1 in the reference A(H3N2) numbering [61].

We used *M* = 360 in the MuE distribution. We set the prior on indels to *μ_r_* = *μ_u_* = (1000, 1) since there is expected to be few indels in this dataset. We trained each model for 7 epochs, which was sufficient for convergence. Hyperparameters and training schedule were otherwise set as in Section S8. To produce the latent embedding in Figure 5, however, we annealed the ELBO weighting χ only up to χ_0_ = 0.001 after 7 epochs, forcing the FactorMuE model into the autoencoding limit (Section S6) [52].

To generate sequences and visualize features, we trained the RegressMuE model on the full time period (1968 to 2019), with 5% of datapoints randomly held out to choose among training runs. To generate future sequences, we sampled from

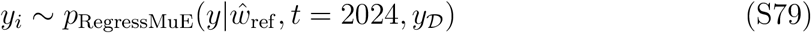

using the variational approximation to the posterior; here 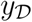 is the training dataset and 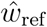 is the maximum *a posteriori* estimator of *w* for the sequence *y*_ref_ of the reference protein structure PDB:4O5N (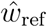 is estimated using a single sample from the variational approximation to the posterior and the Viterbi algorithm). We computed the magnitude of the shift in sequence space from time *t*_0_ to time *t*_1_ in the RegressMuE as

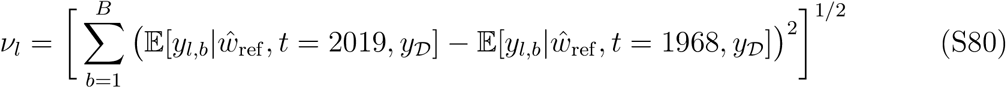

The expectation is estimated using the variational approximation to the posterior with 10 Monte Carlo samples. In evaluating the association between the shift vector *ν*_*l*_ and epitope regions of HA1, we specifically compared to the 16 sites with clear antigenic selection in at least one human sera identified in Lee et al. [28].

## Footnotes

1 Will be made available during peer review at https://github.com/debbiemarkslab.

